# SMARCD1 is a “Goldilocks” metastasis modifier

**DOI:** 10.1101/2024.01.24.577061

**Authors:** Christina Ross, Li-Yun Gong, Lisa M. Jenkins, Ngoc-han Ha, Megan Majocha, Kent Hunter

**Affiliations:** Laboratory of Cancer Biology and Genetics, Metastasis Susceptibility Section, Center for Cancer Research, National Cancer Institute, Bethesda, Maryland, United States of America; Guangdong Provincial Key Laboratory for Genome Stability and Disease Prevention, Department of Biochemistry and Molecular Biology, School of Basic Medical Sciences, Health Science Center, Shenzhen University, 518060, Shenzhen, Guangdong, P. R. China; Laboratory of Cell Biology, Mass Spectrometry Resource, Center for Cancer Research, National Cancer Institute, Bethesda, Maryland, United States of America

## Abstract

Breast cancer is the most frequently diagnosed cancer worldwide, constituting around 15% of all diagnosed cancers in 2023. The predominant cause of breast cancer-related mortality is metastasis to distant essential organs, and a lack of metastasis-targeted therapies perpetuates dismal outcomes for late-stage patients. However, through our use of meiotic genetics to study inherited transcriptional network regulation, we have identified a new class of “Goldilocks” genes that are promising candidates for the development of metastasis-targeted therapeutics. Building upon previous work that implicated the CCR4-NOT RNA deadenylase complex in metastasis, we now demonstrate that the RNA-binding proteins (RNA-BPs) NANOS1, PUM2, and CPSF4 also regulate metastatic potential. Using cell lines, 3D culture, mouse models, and clinical data, we pinpoint *Smarcd1* mRNA as a key target of all three RNA-BPs. Strikingly, both high and low expression of *Smarcd1* is associated with positive clinical outcomes, while intermediate expression significantly reduces the probability of survival. Applying the theory of “essential genes” from evolution, we identify an additional 50 genes that span several cellular processes and must be maintained within a discrete window of expression for metastasis to occur. In the case of *Smarcd1*, small perturbations in its expression level significantly reduce metastasis in laboratory mouse models and alter splicing programs relevant to the ER+/HER2-enriched breast cancer subtype. The identification of subtype-specific “Goldilocks” metastasis modifier genes introduces a new class of genes and potential catalogue of novel targets that, when therapeutically “nudged” in either direction, may significantly improve late-stage patient outcomes.

## Introduction

Metastatic breast cancer remains the leading cause of cancer-related death among women world-wide^1^. In the United States, breast cancer is estimated to be the most highly diagnosed cancer of 2023 and the fourth highest cause of cancer-related mortality after lung, colorectal, and pancreatic cancers. While non-metastatic breast cancer has a 5-year survival rate of 99.3%, metastatic disease reduces the 5-year survival rate to only 31% ^2,3^. For decades, clinicians have targeted cancers by exploiting the unique characteristics that distinguish transformed cells from normal ones, as exemplified by the use of cytotoxic chemotherapy, which takes aim at the hyper-proliferative quality of tumor cells. More recently, genes whose aberrant expression or activity are associated with patient survival have been successfully targeted therapeutically, significantly reducing toxicity and improving the survival of specific patient groups^4^. However, due to the highly evolved nature of metastatic tumors, likely induced by adaptations necessary to complete the metastatic cascade, late-stage patients often do not respond effectively to targeted therapies. Therefore, to improve outcomes for patients with advanced breast cancer, specific metastasis-targeted strategies must be developed, along with a deeper understanding of the unique biological processes that occur during disease progression^5^.

Unfortunately, our understanding of the origins of metastasis is less advanced than that of primary tumorigenesis. This is partially due to the lack of suitable tissue samples, as metastases are often not surgically resected. In addition, most metastatic tissue samples are confounded by prior treatment, heightening the challenge of distinguishing events causing metastasis from those associated with acquired resistance. Moreover, recent sequencing studies from both patients and mouse models suggest that there are no high frequency, commonly mutated metastasis driver genes analogous to tumor drivers^6–8^. The lack of metastasis-specific constitutive activating or inactivation mutations implies that metastasis is more likely driven by transcriptional plasticity directed by epigenetic and microenvironmental signals, consistent with recent studies^9^. Despite considerable progress in this field, much of the etiology of metastasis remains unknown and challenging to identify due to the absence of somatically acquired mutational “fingerprints” that would otherwise highlight crucial components of the metastatic cascade.

An alternative strategy for identifying metastasis-associated genes involves conducting meiotic screens to uncover inherited metastasis susceptibility genes, akin to human epidemiology studies. The identification of candidate genes through this unbiased approach offers an opportunity to explore genes and pathways that might not have been expected to contribute to metastatic progression. To implement this strategy, we have previously utilized the highly metastatic breast cancer MMTV-PyMT mouse model in multiple mouse genetic mapping studies to identify metastasis-associated polymorphic regions of the mouse genome. Through integration of this data with human population genetics, we have described several polymorphism-associated genes that drive metastatic susceptibility in patients^10–14^. This approach has yielded a growing list of candidate metastasis susceptibility and tumor progression genes with potential tumor-autonomous and/or stromal effects, many of which stratify patient outcomes when differentially expressed and hold potential as actionable clinical targets. Importantly, most of the metastasis susceptibility genes identified to date have not been previously implicated in tumorigenesis or metastasis, illustrating the utility of this strategy in uncovering novel pathways and mechanisms in tumor progression.

One of our previous studies implicated the CCR4-NOT RNA deadenylase complex as an inherited breast cancer metastasis factor^15^. Given that the CCR4-NOT complex is a non-specific enzyme, we focused on NANOS1, PUM2 and CPSF4, sequence-specific RNA binding proteins (RNABPs) that were identified in a CCR4-NOT-associated transcriptional network, to gain a better understanding of its role in metastatic disease. These RNA binding proteins recruit specific transcripts to RNA degradation complexes, thereby having the potential to alter molecular pathways through the degradation of specific mRNAs^16–18^. In this study, we demonstrated that NANOS1, PUM2, and CPSF4 function as metastasis-associated factors through post-transcriptional regulation of the SWI/SNF complex component, *Smarcd1*. Unexpectedly, unlike previously identified metastasis susceptibility genes, *Smarcd1* does not follow a linear relationship with patient survival. Instead, we observed a more complex “Goldilocks” effect, where both low and high expression reduce metastasis and improve survival, but an intermediate level is associated with worse outcome. Examination of other SWI/SNF components and components of other molecular complexes suggests that this “Goldilocks” effect may not be limited to *Smarcd1*. If true, this suggests that there may be a set of genes whose activity is narrowly constrained during the metastatic cascade. Targeting these genes to either increase or decrease activity beyond the “Goldilocks” limits may, therefore, provide an additional clinical strategy for the prevention or treatment of metastatic disease.

## Results

### RNA binding proteins NANOS1, PUM2, and CPSF4 modify metastasis

We previously demonstrated that both the structural and catalytic subunits constituting the CCR4-NOT mRNA deadenylation complex modify metastatic propensity^12^. Transcripts alternatively regulated by the CCR4-NOT catalytic subunit CNOT7 in metastatic mouse mammary cancer cell lines were highly enriched for mRNAs containing the canonical binding elements for the RNA-binding proteins (RNA-BPs) NANOS1, PUMILLIO2 (PUM2), and cleavage and polyadenylation specific factor 4 (CPSF4) (Fig. 1a). Furthermore, these transcripts were significantly associated with breast cancer patient outcomes^12^. Based on these findings, we hypothesized that NANOS1, PUM2, and CPSF4 may also serve as metastasis modifiers in breast cancer cells. Indeed, Kaplan-Meier (KM) analysis of breast cancer patients grouped by subtype and using median expression of *NANOS1*, *PUM2*, AND *CPFS4* as a signature, revealed significant stratification of distant metastasis-free survival (DMFS) in patients with the basal subtype (Fig. 1b).

**Figure 1.**
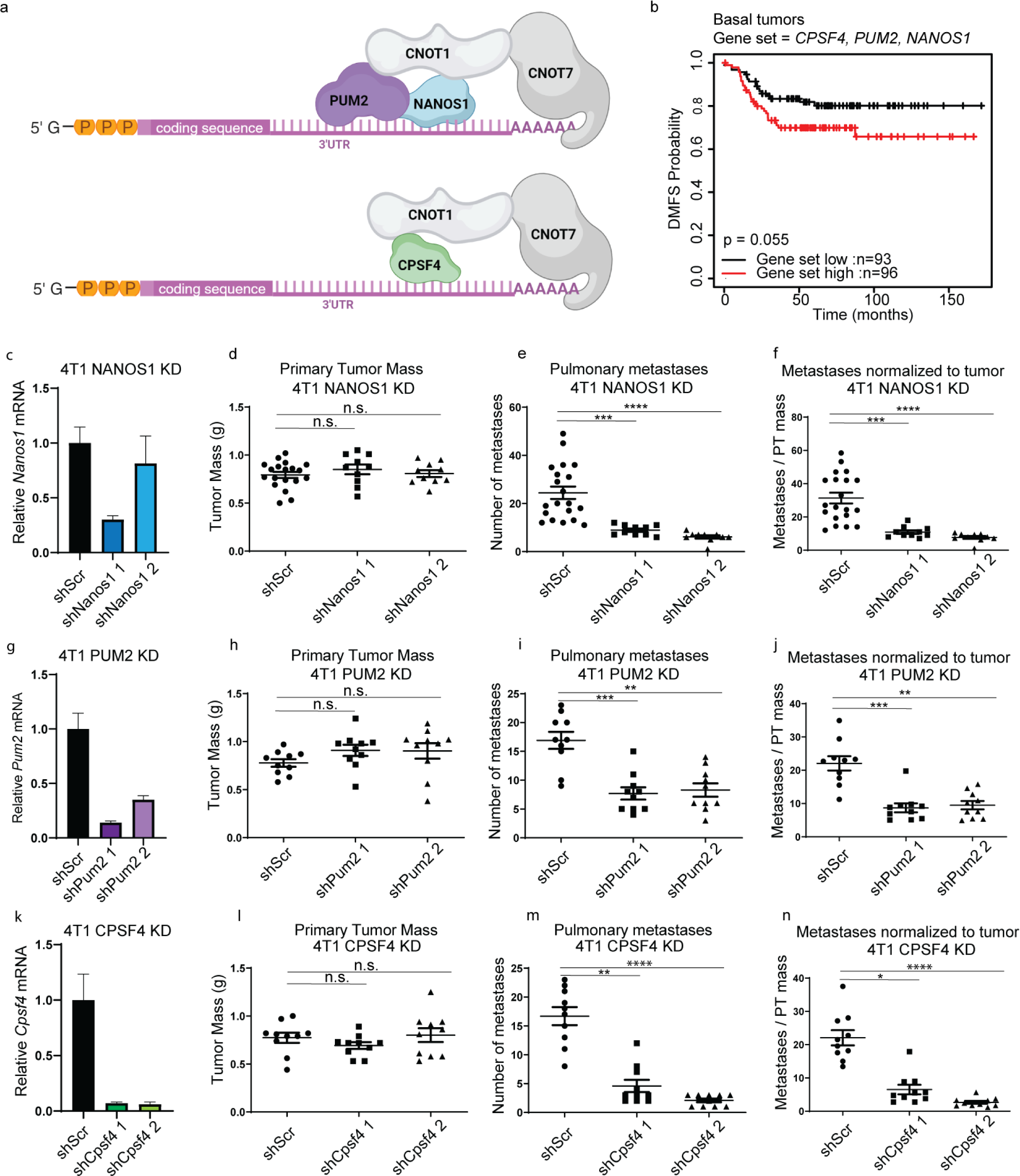
RNA-binding proteins NANOS1, PUM2, and CPSF4 are metastasis modifiers in mouse mammary breast cancer cells. Figure 1 a, representative diagram depicting interactions between mRNA, the deadenylase complex, and RNA-binding proteins PUM2, NANOS1, and CPSF4. b, Kaplan-Meier distant metastasis-free survival (DMFS) for the breast cancer Basal subtype stratified by *PUM2*, *NANOS1*, and *CPSF4* unweighted expression signature in the GOBO database. c, qRT-PCR analysis of *Nanos1* RNA level in 4T1 shScr, or shNanos1 or 2 cells. d-f, analysis of primary tumor (PT) weight (d), number of lung nodules (e), and number of lung nodules per gram of PT per mouse (f) 28 days after orthotopic injection of 4T1 shScr or shNanos1 KD lines. g, qRT-PCR analysis of *Pum2* RNA level in 4T1 shScr, or shPum2-1 or 2 cells. h-j, analysis of PT weight (h), number of lung nodules (i), and number of lung nodules per gram of PT per mouse (j) 28 days after orthotopic injection of 4T1 shScr or shPum2 KD lines. k, qRT-PCR analysis of *Cpsf4* RNA level in 4T1 shScr, or shCpsf4-1 or 2 cells. l-n, analysis of PT weight (l), number of lung nodules (m), and number of lung nodules per gram of PT per mouse (n) 28 days after orthotopic injection of 4T1 shScr or shCpsf4 KD lines. n.s. = not statistically significant, * = p<0.05, ** = p<0.01, ***=p<0.001, ****=p<0.0001.

To more directly assess the role of NANOS1, PUM2, and CPSF4 in metastasis, we knocked down (KD) their expression in two metastatic mouse mammary cancer cell lines, 4T1 and 6DT1, using short hairpin RNAs (shRNAs) (Fig. S1a-b) and performed spontaneous metastasis assays by mammary fat pad injection into syngeneic (BALB/c for 4T1, FVB/N for 6DT1) mice. In both 6DT1 and 4T1 cell lines, KD of NANOS1, PUM2, and CPSF4 did not consistently affect primary tumor growth but did significantly reduce the number of metastatic nodules on the lungs compared to shScramble control (shScr) (Fig. 1c-n and Fig. S1c-n). These data suggest that NANOS1, PUM2, AND CPSF4 act as mediators of breast cancer metastasis.

### NANOS1, PUM2, and CPSF4 regulate the mRNA half-life of SWI/SNF protein SMARCD1

NANOS1, PUM2, and CPSF4 function as key regulators of gene expression by binding to *cis* regulatory elements within mRNAs to recruit the deadenylase machinery^16–18^. Consequently, we performed RNA-sequencing (RNA-seq) to identify gene transcripts altered upon KD of these three factors (Fig. 2a). In 4T1 NANOS1 KD lines, 917 transcripts were altered more than 1.5-fold compared to the control (Fig. 2b, Table S1). Ingenuity pathway analysis (IPA) revealed that these transcripts encoded factors important for cell cycle, DNA damage repair, and several biosynthesis pathways (Table S2). In 4T1 PUM2 KD lines, only 58 transcripts were significantly altered more than 1.5-fold compared to the control (Fig. 2b). These transcripts encoded proteins necessary for several biosynthesis and metabolism programs (Table S2). Finally, in 4T1 CPSF4 KD lines, 1931 transcripts were altered more than 1.5-fold, encompassing pathways related to cell cycle control, DNA damage response, and metabolism (Figure 2b and Table S2). Additionally, when grouping patients by subtype and using each gene list as a signature for KM analysis, NANOS1-, PUM2-, and CPSF4-responsive genes stratified DMFS for patients with the HER2-enriched subtype of breast cancer (Fig. S2a-c). Venn diagram analysis of the gene lists identified 21 transcripts with differential expression in all 4T1 KD lines compared to their matched control (Fig. 2b-d and Table S1). Using functional annotation clustering and the database for annotation, visualization and integrated discovery (DAVID), these 21 factors were categorized into 4 major groups: 1) cell morphogenesis, 2) nuclear lamina, 3) cell motion and adhesion, and 4) extracellular signaling (Table S1). When used as a 21-gene expression signature, KM analysis showed significant stratification of breast cancer patient DMFS (Fig. 2e). To complement the 4T1 RNA-seq, we performed qRT-PCR for 6DT1 KD and control lines. Nine consistently altered transcripts were identified (Fig. 2f-h) and screened as direct targets of NANOS1, PUM2, and CPSF4.

**Figure 2.**
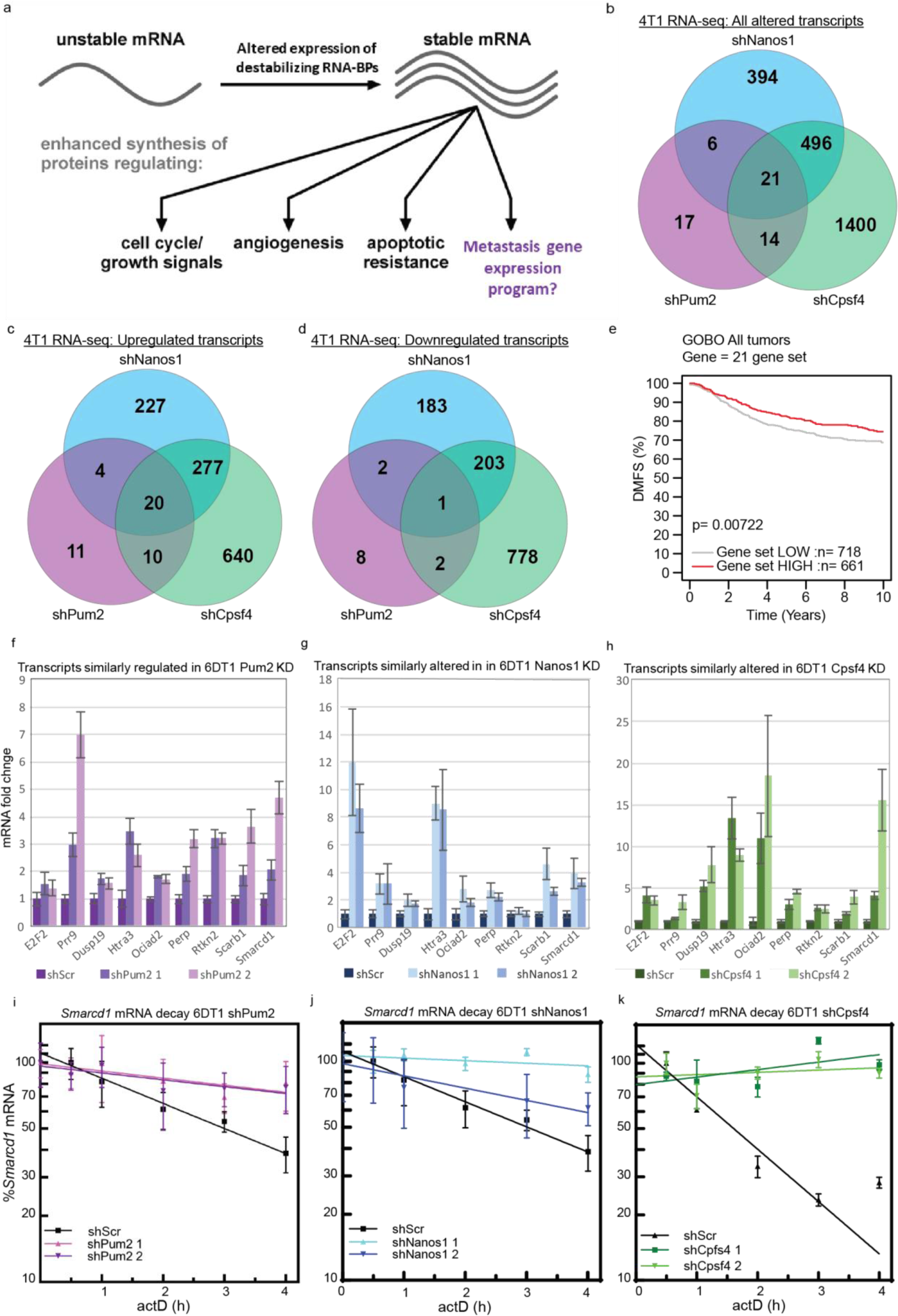
NANOS1, PUM2, and CPSF4 regulate *Smarcd1* mRNA half-life. Figure 2a, diagram illustrating a hypothetical metastatic gene expression program regulated by RNA-binding protein (RNA-BP) abundance and mRNA stability. b, Venn-diagram analysis of RNA-seq differential gene expression (≥1.5 fold change ≤-1.5) and false discovery rate (FDR) less than 0.05 from 4T1 shNanos1 vs shScr, shPum2 vs shScr, and shCpsf4 vs shScr. c-d, Venn-diagram analysis of RNA-seq differential gene expression for genes with ≥1.5 fold change (c) and ≤-1.5 fold change (d) with FDR of 0.05. e, Kaplan-Meier analysis of distant metastasis-free survival (DMFS) in all breast cancer patients stratified by 21 commonly differentially expressed genes (used as a non-weighted signature) in the GOBO database. f-h, nine genes with consistent expression changes by qRT-PCR analysis of 21 differentially expressed genes in 6DT1 shPum2 (f), shNanos1 (g), and shCpsf4 (h) vs shScr (n=3). i-k, representative actinomycin D (actD) time courses showing *Smarcd1* transcript decay in 6DT1 shPum2 (i), shNanos1 (j), and shCpsf4 (k) vs shScr lines (n=3).

To determine if the nine commonly altered transcripts were directly regulated by NANOS1, PUM2, or CPSF4, we manually surveyed the 3’ untranslated regions (3’UTRs) of the mature RNA (mRNA) transcripts for canonical *cis*-response elements^16,17,19,20^. *Smarcd1*, *Prr9*, and *E2f2* mRNA contained a PUM2 recognition element (PRE), partial NANOS RE (NRE), and Cpsf4 recognition sequence (Fig. S3a-d). To determine if the *Smarcd1*, *Prr9*, or *E2f2* mRNA half-life was dependent on NANOS1, PUM2, and CPSF4 levels, we performed actinomycin D (actD) time courses in 6DT1 control and KD cell lines. *E2f2* mRNA decay was not altered by the KD of NANOS1, PUM2, or CPSF4 (Fig. S3e and Table S3), while both *Prr9* and *Smarcd1* transcripts were significantly stabilized in the KD cell lines (Fig. 2i-k, Fig. S3f and Table S3). *Prr9* mRNA half-life was approximately 3.4 hrs in 6DT1 control cells, and this was extended to over 6 hrs in the KD lines (Fig. S3f and Table S3). Similarly, *Smarcd1* mRNA half-life was approximately 2.9 hrs in 6DT1 control cells and over 6 hrs in the KD lines (Fig. 2i-k and Table S3). These results support the hypothesis that *Prr9* and *Smarcd1* mRNAs are direct targets for destabilization by NANOS1, PUM2, and CPSF4.

Interestingly, *Smarcd1* is a member of the “Role of BRCA1 in DNA Damage Response” pathway, which was the most significantly enriched pathway according to IPA of differential gene expression in PUM2 and CPSF4 KD cells, and the fourth most significantly altered pathway in NANOS1 KD cells (Table S2). Additionally, *Smarcd1* can be found within the “Cell Morphogenesis” functional annotation group, which was the only significantly populated group identified by DAVID analysis of the 21 commonly dysregulated genes (Table S1). Based on these findings, we hypothesized that the direct regulation of the *Smarcd1* transcript by NANOS1, PUM2, and CPSF4 may be necessary for the metastasis of breast cancer cells.

### Dysregulation of *Smarcd1* increases tumorsphere formation

To test if *Smarcd1* expression could modify cellular processes associated with breast cancer progression *in vitro,* we created 6DT1 cells with knockdown (KD) or overexpression (OE) of *Smarcd1*, as well as appropriate control lines shScr or empty vector (EV), respectively (Fig. 3a). *Smarcd1* OE and KD did not impact cell proliferation under normal growth conditions or upon glutamine deprivation, exposure to increased reactive oxygen species, heat shock, or hypoxia (Fig. S4a-d). Similarly, cell sensitivity to the DNA-damaging agent doxorubicin (Dox) or continuous heat shock stress was not altered by *Smarcd1* levels (Fig. S4e-f). Furthermore, 2D colony formation assays and cell migration (scratch assays) did not reveal any differences in phenotype dependent on *Smarcd1* expression (Fig. S4g-i).

**Figure 3.**
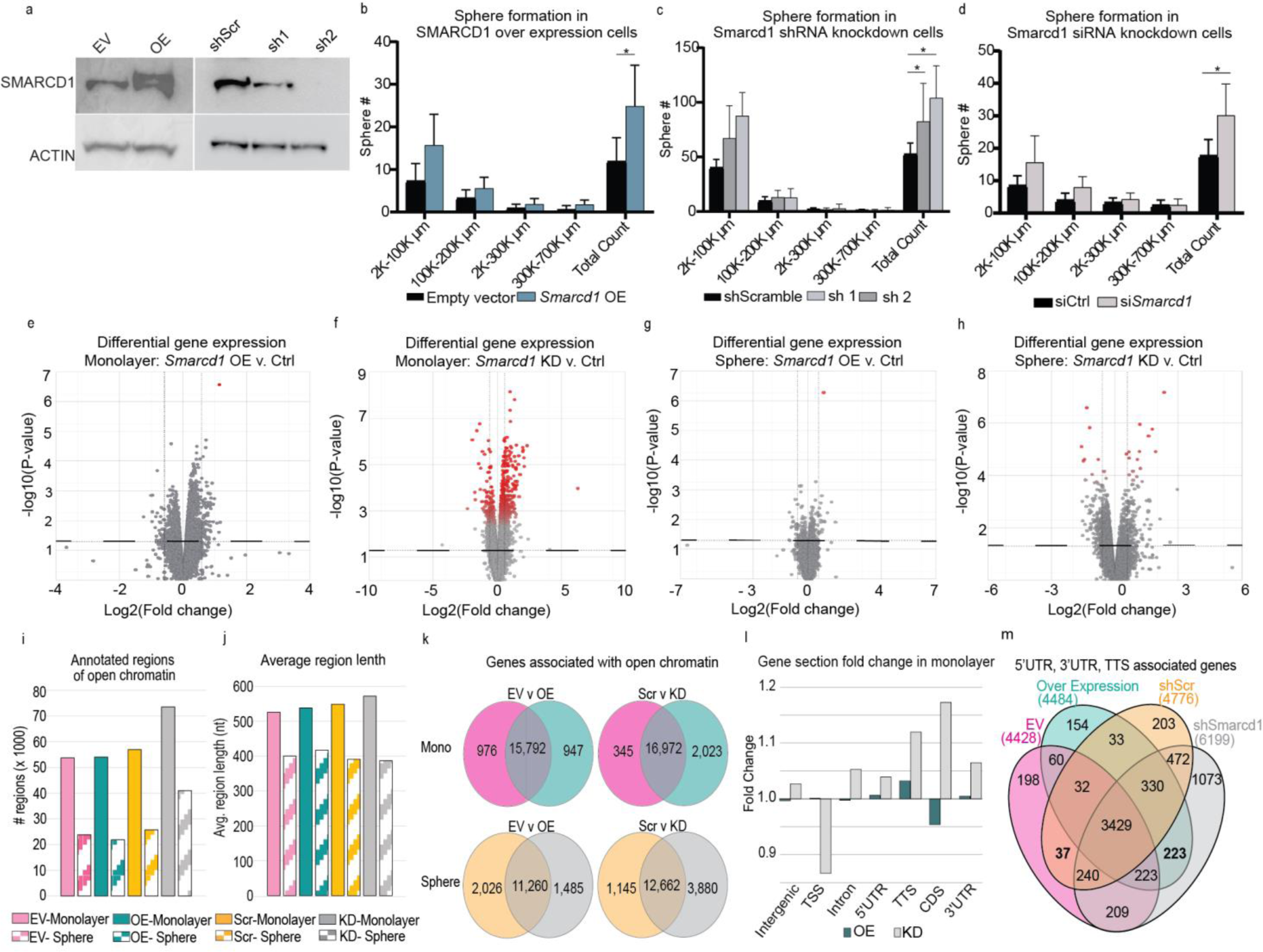
*Smarcd1* expression regulates tumorsphere formation and chromatin accessibility. Figure 3a, Western blot showing SMARCD1 and Actin protein levels after transduction of 6DT1 cells with empty vector (EV), *Smarcd*1 overexpression vector (OE), shScramble (shScr), and shSmarcd1 shRNA-1 and shRNA-2. b-d, tumorsphere number grouped by size 10 days after seeding 6DT1 EV control and *Smarcd1* OE lines (b), 6DT1 shScr control and shSmarcd1 lines (c), or 6DT1 cells transfected with *Smarcd1*-targeted siRNA or non-targeting control (d), representative data from 1 of 3 replicates, n=6 wells per cell line. * = p<0.05. e-h, volcano plots showing differential gene expression analysis with false discovery rate (FDR) < 0.1 indicated by red intensity in *Smarcd1*-altered 6DT1 cell lines vs controls, in monolayer culture (e-f) and sphere culture (g-h). i-j, histogram showing number of regions of open chromatin (i) detected by ATAC-seq analysis and their average length (j). k, Venn diagram analysis of the genes associated with open regions of chromatin in *Smarcd1*-altered cells vs controls and in monolayer culture and in sphere culture. l, fold change in the enrichment of specific gene sections (Intergenic, Transcription start site or TSS, Intron, 5 prime untranslated region or 5’UTR, Transcription termination sequence or TTS, coding sequence or CDS, and 3’UTR) in open chromatin for *Smarcd1*-altered cells vs control lines grown in monolayer. m, Venn diagram analysis of genes from *Smarcd1*-altered cells and controls grown in monolayer for which 3’ UTR, 5’ UTR, and TTS were enriched in open chromatin.

Unlike 2D culture, cells seeded in methylcellulose for 3D culture formed significantly more spheres, surprisingly for both *Smarcd1* KD and OE cells compared to controls (Fig. 3b-c). However, the sphere size distribution was not significantly altered, suggesting that this phenotype may be related to stemness or anchorage rather than proliferation. This result was further confirmed using siRNA KD (Fig. 3d, Fig. S4j).

### SMARCD1-dependent changes in chromatin accessibility are subtly dependent on culture type

SMARCD1 functions within the chromatin-remodeling BAF complex and has been described as a key mediator of cardiac and myeloid development^21,22^. Thus, we next asked if altered *Smarcd1* expression could modify gene expression programs by changing the regions of open chromatin, and whether any patterns may be common to both *Smarcd1* KD and *Smarcd1* OE cells.

First, RNA-seq was performed with 6DT1 *Smarcd1* KD and OE cells grown in monolayer and 3D culture. Neither *Smarcd1* (Fig. S5a) nor the expression of BAF complex factors was significantly affected by culture type (Fig. S5b-c). Using differential gene expression analysis and a p value cut-off of 0.05, we identified 1,746 genes in *Smarcd1* OE monolayer cells and 2,351 genes in *Smarcd1* KD monolayer cells with altered expression compared to the appropriate control (Fig. 3e-f and Table S4). Fewer genes were altered in sphere culture, with 281 genes in *Smarcd1* OE cells and 1190 in KD cells (Fig. 3g-h and Table S4). Most of the gene expression changes were below 1.5-fold in both culture conditions. Importantly, when we applied a false discovery rate (FDR) statistical test cutoff of 0.1 in addition to a p value cutoff, the number of differentially expressed genes was reduced to only 5 and 1 (*Smarcd1*) genes in *Smarcd1* OE monolayer and sphere cultures, respectively, and 641 and 34 genes in *Smarcd1* KD monolayer and sphere cultures, respectively (Fig. 3e-h, red dots and Table S4).

Despite the minor effects observed on gene expression, we next assessed if *Smarcd1* expression might modulate chromatin structural changes in breast cancer cells. Assay for transposase-accessible chromatin-sequencing (ATAC-seq) was performed with 6DT1 *Smarcd1* KD and OE cells grown in monolayer and sphere culture. Both the number and length of annotated open chromatin regions were reduced in cells from sphere culture compared to monolayer (Fig. 3i-j and Table S5). Additionally, *Smarcd1* KD resulted in more regions of open chromatin in both monolayer and sphere culture compared to controls and OE cells (Fig. 3i). Consistent with this observation and the gene expression data, the number of genes associated with open chromatin was also decreased in sphere vs monolayer samples and in *Smarcd1* OE vs KD (Fig. 3k). Venn diagram analysis of monolayer cells revealed 265 genes associated with open chromatin in *Smarcd1* OE and KD lines that were not accessible in controls, and 35 genes associated with open chromatin in the controls alone and therefore inaccessible in *Smarcd1*-altered lines (Fig. S5d). In sphere culture, 436 genes were associated with open chromatin in *Smarcd1* OE and KD lines, and 158 genes associated with open chromatin in controls (Fig. S5d). Comparison between monolayer and sphere cultures revealed that dysregulation of *Smarcd1* impacted disparate regions of chromatin in a culture type-dependent manner, as only 11 genes were commonly associated with open chromatin in all *Smarcd1*-altered cells (Fig. S5d).

We hypothesized that *Smarcd1* may impact chromatin accessibility at specific gene regions. Gene section analysis revealed that the overall enrichment of each gene section was similar between samples (Fig. S5e and Table S5). Fold-change for the enrichment of specific gene sections in open chromatin was assessed for *Smarcd1*-altered cells compared to controls for monolayer and sphere culture. In sphere culture, *Smarcd1* OE and KD resulted in opposing fold change for each gene section (negative vs positive fold change) compared to controls (Fig. S5f). However, monolayer cells revealed a positive fold change in the enrichment of transcription termination sequences (TTS), 3’ UTR, and 5’UTR gene sections in both *Smarcd1* OE and KD lines (Fig. 3l), with TTS having the largest enrichment. Venn diagram analysis of genes associated with those TTS, 3’ and 5’UTRs identified 223 genes common to open chromatin in *Smarcd1* OE and KD lines (Fig. 3m). However, assessment of these genes in the RNA-seq dataset revealed their expression to be largely unchanged (Fig. S5g).

Finally, the ATAC-seq data was analyzed for unique motif sequences that may be enriched in the open chromatin of *Smarcd1*-altered cells. Consistent with the finding that sphere cultures had fewer and shorter regions of open chromatin, we also found fewer significantly enriched motifs in sphere-derived samples (Fig. S5h and Table S6). By Venn diagram analysis, there were no enriched motifs common to *Smarcd1*-altered cells that were not also observed in control lines for cells grown in monolayer. In sphere culture, two motifs were commonly enriched in the open chromatin of *Smarcd1* OE and KD cells, and three were exclusive to the controls (Fig. S5i and Table S6). One of the enriched motifs was the binding site for retinoic acid receptor alpha (RARA), which has been previously shown to have clinical relevance for breast cancer patients with ER+ tumors^23^.

### SMARCD1 interacts with splicing machinery to regulate clinically relevant splicing programs

The lack of overt changes to gene expression or chromatin structure in *Smarcd1* OE cells was intriguing given the known function of SMARCD1 within the BAF complex and the modification of sphere formation in 3D culture (Fig. 3b-c). To investigate the role of SMARCD1 in metastatic breast cancer cells, we performed co-immunoprecipitation followed by proteomic analysis of lysates from 6DT1 *Smarcd1* OE cells grown in monolayer culture (Fig. S6a-b). Using an endogenous antibody to Smarcd1 protein, enrichment of interacting proteins was calculated in OE cells compared to EV control. Consistent with literature, we identified core BAF complex proteins as well as GBAF, nBAF, and npBAF factors as *Smarcd1*-interactors and these were validated by IP and western blot (Fig. S6c-d). Proteins with fold change in enrichment greater than 5 as compared with a control IP were considered biologically relevant interactors for breast cancer metastasis, and this resulted in a list of 286 factors (Table S7). To identify cellular processes involving SMARCD1, the list was entered into the DAVID tool for functional annotation clustering. The top three enriched clusters were “Chromatin remodeling”, “mRNA processing”, and “SWI/SNF or BAF complex” (Fig. 4a). Additionally, functional classification of the top interactors with a fold change over 1000 was performed manually, and those with known cellular functions were grouped (Fig. 4b). Cellular processes associated with the most highly enriched proteins upheld previously reported canonical BAF interactions such as chromatin remodeling and nuclear lamina but also unexpectedly implicated SMARCD1 in metabolism, exocytosis, RNA regulation, and splicing.

**Figure 4.**
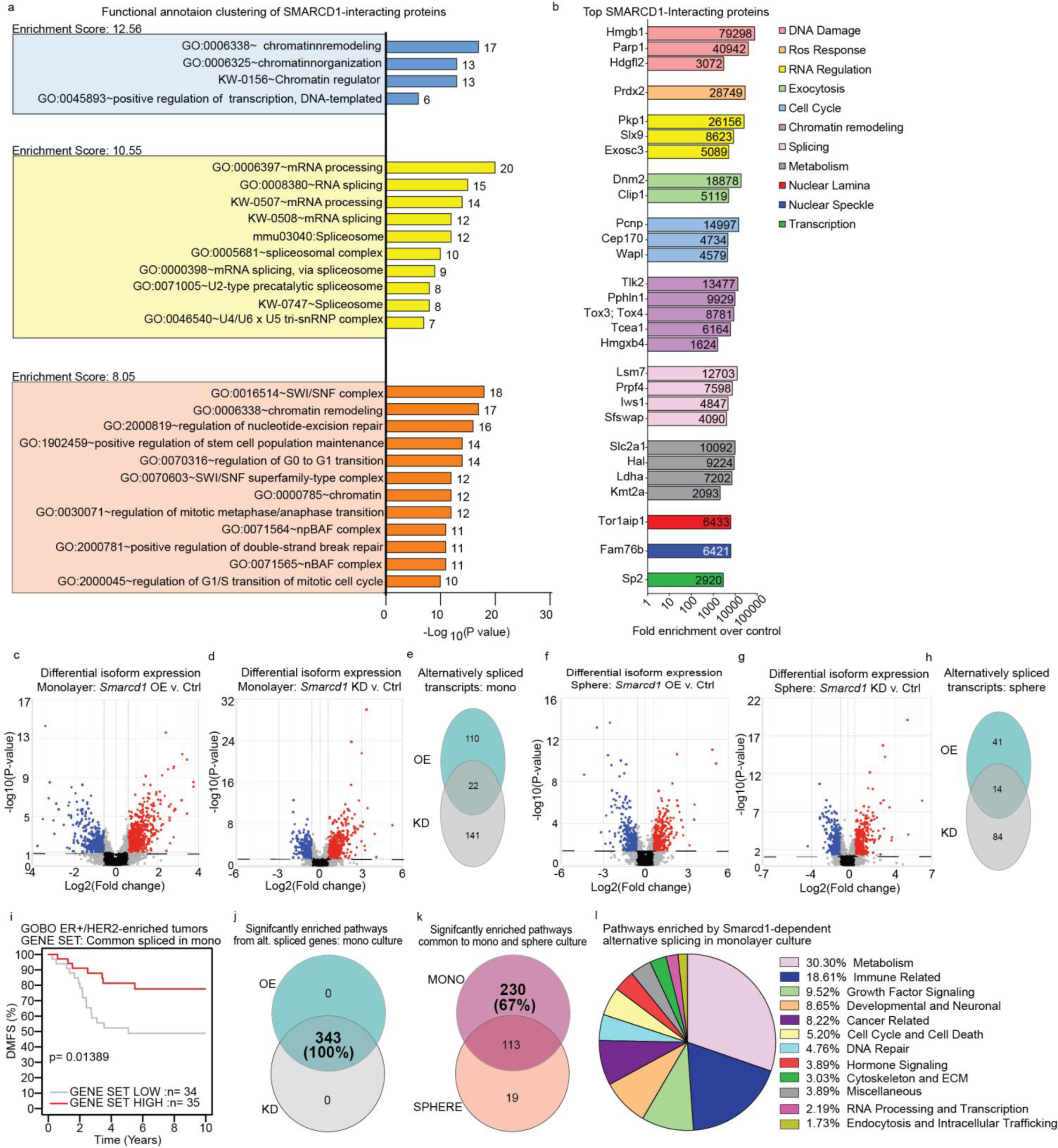
SMARCD1 interacts with splicing machinery and alters a clinically relevant splicing program. Figure 4a, graph of top 3 DAVID functional annotation clusters, enrichment scores, and −Log10(p value) for all SMARCD1-interacting proteins in 6DT1 *Smarcd1* OE cells. b, graph showing functional annotations and fold enrichment for individual SMARCD1-interacting proteins with highest enrichment. c-e, differential isoform expression in 6DT1 *Smarcd1*-altered cells vs controls in monolayer shown on volcano plots (blue: negative fold change, red: positive fold change) for overexpression (OE) vs control (c) and knockdown (KD) vs control (d), with Venn diagram analysis of genes with p<0.05 and false discovery rate (FDR) <0.05 (e). f-h, differential isoform expression in *Smarcd1*-altered cells vs controls in sphere culture shown on volcano plots (blue: negative fold change, red: positive fold change) for OE vs control (f) and KD vs control (g), with Venn diagram analysis of genes with p<0.05 and FDR <0.05 (h). i, GOBO database distant metastasis-free survival (DMFS) Kaplan-Meier analysis for the ER+/HER2-enriched subtype using an unweighted gene signature of 22 common alternatively spliced genes in *Smarcd1*-altered cells vs controls. j, Venn diagram analysis of pathways significantly enriched (p<0.05) with alternatively splicing genes in *Smarcd1* OE vs KD cells. k, Venn diagram analysis of pathway significantly enriched (p<0.05) with alternatively splicing genes common to *Smarcd1*-altered cells in monolayer vs sphere culture. l, Pie chart of pathway categories unique to monolayer culture alternative splicing in *Smarcd1*-altered cells.

To determine if SMARCD1 may play a role in RNA splicing in metastatic breast cancer cells, the RNA-seq data was reanalyzed for transcript level differential expression in *Smarcd1* OE and KD lines in monolayer and sphere culture compared to the corresponding controls. Interestingly, both OE and KD of *Smarcd1* in monolayer and sphere culture resulted in significant changes to specific transcript levels compared to controls. When subject to both p-value and FDR cut-offs, 132 and 163 transcripts were differentially expressed in *Smarcd1*-OE and KD monolayer cells, respectively, of which 22 were common (Fig. 4c-e and Table S8). *Smarcd1* OE and KD in sphere culture resulted in 55 and 98 differentially expressed transcripts, of which 14 were common (Fig. 4f-h and Table S8). When used as a signature for subtype-specific KM analysis, transcripts commonly altered in *Smarcd1* OE and KD lines grown in sphere culture were not associated with patient outcomes. However, the 22 commonly altered transcripts in monolayer culture significantly stratified ER+/HER2-enriched subtype DMFS (Fig. 4i), suggesting an important role for their regulation in this patient group. Pathway analysis was then performed using IPA for *Smarcd1* OE and KD alternatively spliced gene lists for both monolayer and sphere cultures. Unexpectedly, when pathway analysis was visualized by p value cut-off only, pathways enriched by alternatively spliced transcripts in *Smarcd1* OE and KD overlapped completely, and this was observed for both monolayer culture and sphere culture samples (Fig. 4j, Fig. S6e-g and Table S9). As alternative splicing in monolayer culture was significantly associated with patient outcomes (Fig. 4i), those pathways uniquely enriched in monoculture were manually categorized into 11 major cellular processes (Fig. 4k-l). The largest category of enriched pathways was metabolism, followed by immune-related signaling and growth factor signaling, suggesting that these cellular processes may be key for the process of metastasis in the ER+/HER2-enriched breast cancer subtype (Fig. 4l).

### SMARCD1 functions as a “Goldilocks” metastasis modifier

Given that positive and negative modulation of *Smarcd1* expression in mouse mammary cancer cell lines resulted in increased tumorsphere formation and alternative splicing of a common clinically relevant gene set, we next sought to determine if changes in *Smarcd1* expression modify metastasis *in vivo*. Accordingly, 6DT1 *Smarcd1*-altered cells and controls were injected into the mammary fat pad of syngeneic mice to assess their propensity for spontaneous metastasis. We observed that neither OE nor KD of *Smarcd1* significantly altered primary tumor growth. However, both high and low levels of *Smarcd1* expression significantly reduced the number of metastatic nodules on the lungs (Fig. 5a-f and Fig. S7a-c), suggesting that tight regulation of *Smarcd1* expression facilitates breast cancer metastasis in mice. While this pattern was also observed with tail vein injection assays, the trend was no longer significant, suggesting that *Smarcd1* expression likely impacts early steps in the metastatic cascade such as intravasation into the circulation, and not seeding at the lung. Finally, we mined our previously published RNA-seq data of matched primary and metastatic tumors harvested from the Polyoma Middle T genetically engineered mouse model crossed to 8 mouse strains^24,25^. When comparing primary and secondary tumors, there was no difference in *Smarcd1* expression (Fig. S8a), consistent with the hypothesis that elevation or reduction of expression may decrease metastatic propensity.

**Figure 5.**
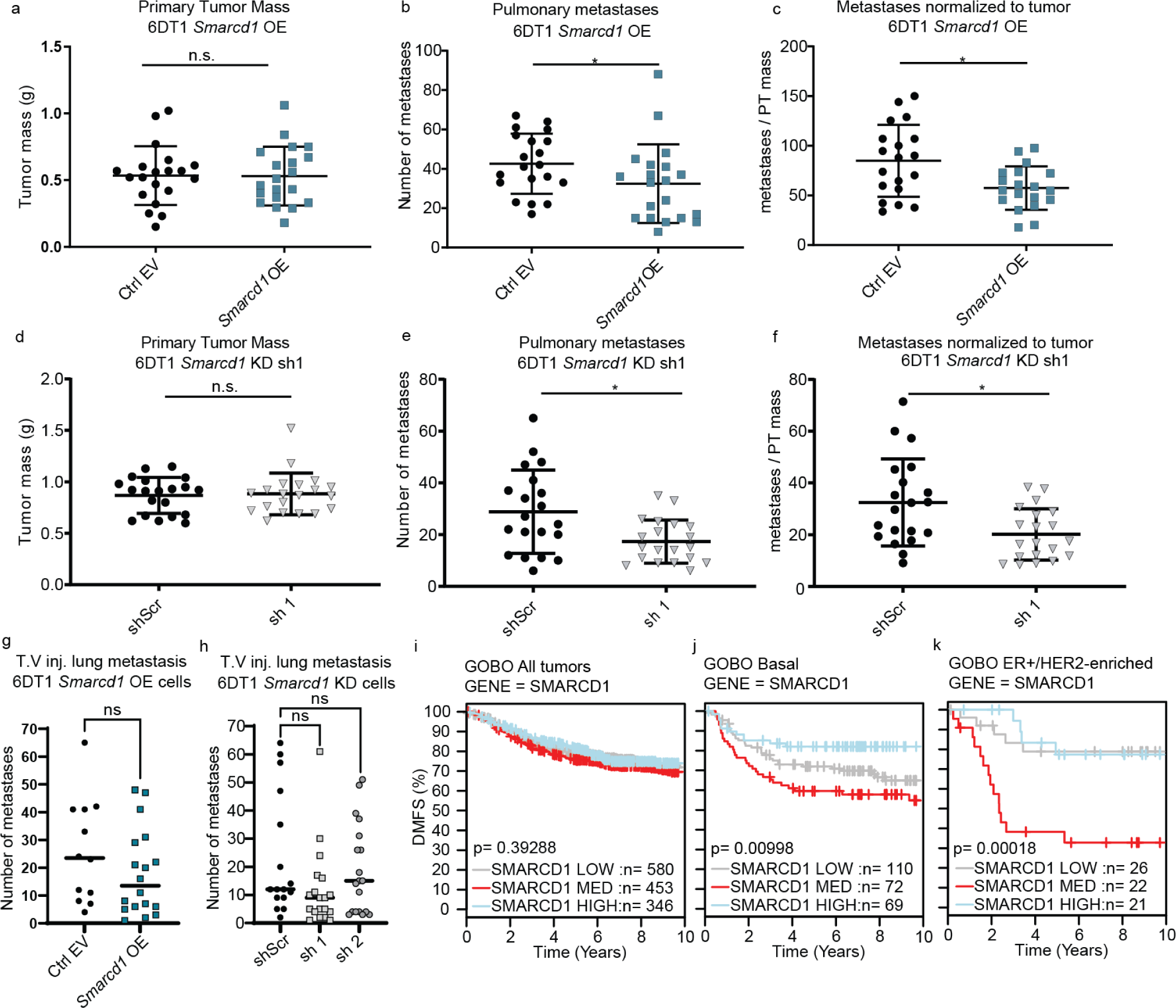
SMARCD1 is a “Goldilocks” metastasis modifier. a-c, analysis of primary tumor (PT) weight (a), number of lung nodules (b), and number of lung nodules per gram of PT per mouse (c) 28 days after orthotopic injection of 6DT1 EV control and *Smarcd1* OE lines (2 combined experiments, 10 mice per experiment). d-f, analysis of PT weight (d), number of lung nodules (e), and number of lung nodules per gram of PT per mouse (f) 28 days after orthotopic injection of 6DT1 shScr control and shSmarcd1 lines (2 combined experiments, 10 mice per experiment). g-h, number of lung metastases per mouse 28 days after tail vein injection (TV inj) of 6DT1 EV control and *Smarcd1* OE lines (g) or 6DT1 shScr control and shSmarcd1 lines (h) (2 experiments combined, 10 mice per experiment). i-k, GOBO database distant metastasis-free survival (DMFS) Kaplan-Meier analysis stratified by *SMARCD1* expression level for all breast cancer patients (i), Basal subtype patients (j), or ER+/HER2-enriched subtype patients (k).

To determine if SMARCD1 plays a role in the progression of human disease, we queried each breast cancer subtype using the Gene expression-based Outcome for Breast cancer Online (GOBO) dataset^26^. Dysregulation of *SMARCD1* significantly stratified DMFS for Basal- and ER+/HER2-enriched patient subtypes (Fig. 5i-k). Interestingly, and consistent with our *in vivo* data, both high and low *SMARCD1* levels were associated with better outcomes for these subtypes, while “intermediate” *SMARCD1* expression was associated with worse outcome (Fig. 5j-k). We next asked if other genes followed a similar pattern, functioning as “Goldilocks metastasis modifiers”. Through a manual search of well-characterized genes in the GOBO dataset, 29% of factors involved in chromatin remodeling, transcription, RNA processing, and metabolism followed a Goldilocks pattern of DMFS stratification for one or more breast cancer subtype (Fig. S8b and Table S10). A specific analysis of the SWI/SNF complex revealed that, in addition to SMARCD1, Golidlocks survival curves were observed for core the BAF proteins BCL7A, BCL7C, and BRG1 in ER+ patients, accessory nBAF proteins CREST and BRD9 in ER+ Luminal A patients, and accessory esBAF protein BCL11B in Basal subtype patients (Table S10)^27^. Taken together, the data reported here are consistent with the existence of a “Goldilocks” gene group that spans many cellular processes and complexes, for which a tight regulation of intermediate expression is necessary for metastasis within discrete breast cancer subtypes. More specifically, SMARCD1 functions as a Goldilocks modifier in ER+/HER2-enriched breast cancer metastasis through the regulation of splicing.

## Discussion

Tumor progression and metastasis is a complex, systemic process that involves multiple tissues and cell types throughout the body^28^. Evidence suggests that as early tumors become hypoxic, tumor cells co-opt developmental transcriptional programs that enable invasion and motility to escape the toxic environment of the primary tumor bed. Although dissemination can occur before tumors reach the lower limit of clinical detection, early resection of breast cancer lesions is associated with an extremely high 5-year survival rate, suggesting that additional events must occur before disseminated tumor cells can give rise to clinically relevant macroscopic lesions^28^. In contrast to tumor initiation, sequencing studies in both patient and animal models have not identified high-frequency, common metastasis driver mutations. This implies that somatic constitutive activation or inactivation of genes does not play a major role in metastatic progression^6,8^. Instead, current models propose that activation of developmental mechanisms, such as epithelial-to-mesenchymal-like transition or transient acquisition of stem cell-like capacities, are likely critical drivers of metastatic development^9^. Consequently, understanding the regulation and epigenetic integration of tumor-autonomous and microenvironmental interactions related to this cellular plasticity is paramount for deciphering the processes and vulnerabilities that underlie the proximal cause of most breast cancer mortality.

To investigate this, our laboratory adopted a meiotic genetics approach to identify metastasis susceptibility genes, primarily based on inherited variations that alter gene transcription^29,30^. In this study, we expanded upon previous observations that implicated the CCR4-NOT mRNA deadenylation complex as a metastasis susceptibility factor^12^. Given that the CCR4-NOT complex functions as a non-specific deadenylase, we hypothesized that RNA binding partners recruit specific metastasis-associated transcripts for degradation, and variations in the efficiency of this process alter the metastatic capacity of tumor cells. Consistent with this hypothesis, our findings indicate that CCR4-NOT-associated mRNA binding proteins, namely NANOS1, PUM2, and CPSF4, all contribute to metastasis in allograft orthotopic spontaneous metastasis assays. Moreover, the KD of all three mRNA binding proteins resulted in the stabilization the transcript encoding the SWI/SNF complex protein SMARCD1, which we subsequently demonstrated also alters metastatic capacity.

Unexpectedly, in contrast to studies on other metastasis susceptibility genes that revealed simple linear relationships between transcription levels and metastatic capacity, both ectopic expression and shRNA-mediated KD of *Smarcd1* suppressed metastatic capacity compared to intermediate transcription levels. The mechanism by which SMARCD1 influences metastasis is likely associated with its role in transcriptional control through chromatin modification. To complete the metastatic cascade, tumor cells must transiently engage several pathways, including mechanisms necessary to withstand the toxic primary tumor microenvironment, initiating motility and invasion programs, surviving in transit, evading the immune system, and adapting to novel microenvironments at the secondary site^28^. Considering this, the observation that either up- or down-regulation of SMARCD1 resulted in an increase in tumor sphere formation, a feature associated with stem-like characteristics and increased metastatic capacity, was unexpected^31^. In the case of SMARCD1, we hypothesize that increased expression may have “locked” chromatin into a configuration less adaptable to changing metastatic transcriptional requirements. In contrast, cells with reduced SMARCD1 levels exhibited increased chromatin accessibility, which may hinder the ability of cells to orchestrate orderly transcriptional changes due to inappropriate chromatin conformations. Thus, the dysregulation of SMARCD1 may impede the cellular plasticity between stem-like states believed to be essential for metastatic colonization. However, further work will be necessary to address this possibility.

Interestingly, both increased and decreased SMARCD1 levels resulted in differential splicing without significant changes in the overall transcriptional program in bulk cell populations. Several studies have shown that chromatin remodeling complexes can interact with the spliceosome and that splicing factors often associate with nucleosomes, suggesting that dynamic chromatin structure plays an important role in splicing^32,33^. In a recent study by Reddy et al., the BAF complex protein ARID1B was shown to interact with the long non-coding RNA NEAT1 and the paraspeckle to regulate alternative splicing^34^. In this study, we found that SMARCD1 interacts with ARID1B as well as several RNA processing and splicing factors. Therefore, it is possible that the splicing changes we observed may be attributed to an ARID1B-dependent paraspeckle interaction directed by SMARCD1.

The “Goldilocks”-like phenomenon was also observed for *SMARCD1* in some subtypes of human breast cancer, suggesting that this result is not merely an artifact of laboratory model systems. Examination of other members of the SWI/SNF complex and additional cellular complexes suggests that this “Goldilocks” phenotype may extend beyond SMARCD1, as tumor cells cannot tolerate significant alterations in several basic molecular functions while also retaining metastatic capacity. Indeed, our examination of clinical data for 15 biological complexes identified 50 additional “Goldilocks genes” spanning multiple cellular processes. Interestingly, the genes we identified in this class, like *SMARCD1*, exhibit high context dependence, and even individual factors within one molecular complex can stratify disparate clinical subtypes. This implies that different subtypes of breast cancer likely utilize different mechanisms to achieve metastatic competence. This hypothesis aligns with our previous demonstration of subtype specificity for different metastasis susceptibility genes, suggesting the existence of multiple pathways to achieve metastatic colonization.

“Goldilocks” effects, though potentially novel in the setting of tumor evolution, have been described in the context of organismal evolution. Briefly, while the amplification and deletion of many genes can be tolerated and even beneficial for species survival, there exists another group of genes for which change is not tolerated outside of a specific life-defining range^35^. Applying this concept to tumor evolution suggests that there are functions and biological processes that must be maintained at specific levels to enable metastatic progression. Moreover, since different breast cancer subtypes exhibit distinct cellular phenotypes, the molecular characteristics defining each subtype also determine their vulnerabilities and tolerance to change. This, in turn, dictates which molecular processes are restricted to specific viable windows of activity. In the case of the ER+/HER2-enriched subtype, the SMARCD1-directed regulation of splicing programs important for metabolism, immune signaling, and differentiation emerges as a key factor for metastasis.

To the best of our knowledge this is the first time a “Goldilocks” metastasis modifier gene has been reported in the literature. This is not entirely surprising, given that target identification typically involves stratifying data using binary or linear patterns and selecting characteristics that differentiate metastatic tumors from primary tumors and normal tissue. Interestingly, reanalysis of RNA-seq data from a previous PyMT GEMM study revealed no difference in *Smarcd1* expression between matched primary and metastatic tumors across several genetic backgrounds. This lack of difference would typically categorize a gene as clinically irrelevant in the study of metastasis, but it is now evident that a narrow range of expression may be essential for metastatic progression. Therefore, the identification of “Goldilocks” genes may require a divergence from seeking linear relationships. For example, reanalyzing primary and metastatic tumor exome-sequencing data could identify genes with infrequent copy number variation, and KM analysis of patient gene expression datasets could be presented in a tripartite manner for a more nuanced exploration.

Importantly, the existence of these “Goldilocks” genes may open additional avenues for clinical intervention. Current therapeutic development predominantly focuses on generating small molecule or antibody inhibitors designed to interfere with protein function. However, for “Goldilocks”-type proteins, either increased or decreased function may prove efficacious against metastatic disease, particularly if the “Goldilocks” window is relatively narrow. For *SMARCD1*, the up- or down-regulation achieved was relatively modest (∼1.5-fold), which might be more easily attainable than the suppression needed to target oncogenic drivers. Furthermore, the small perturbations that significantly reduced metastasis did not impact cell survival or proliferation, suggesting that therapeutically targeting SMARCD1 may cause less systemic toxicity. Combining “Goldilocks” activation/suppression with more conventional targeted therapies may, therefore, offer an additional and alternative method for adjuvant therapy or targeting established metastatic lesions.

## Materials and Methods

### Cell culture

Mouse mammary carcinoma cell lines 4T1 and 6DT1 were a generous gift from Dr. Lalage Wakefield (NCI, Bethesda, MD). All cell lines were cultured in Dulbecco’s Modified Eagle Medium (DMEM), supplemented with 10% fetal bovine serum (FBS), 1% penicillin and streptomycin (P/S), and 1% glutamate, and maintained at 37°C with 5% CO_2_. Short hairpin RNA (shRNA)-mediated knockdown and overexpression cells were cultured in the same conditions with an addition of 10 µg/ml puromycin and 5 µg/ml blasticidin, respectively.

### Plasmid constructs

#### shRNA constructs

TRC lentiviral shRNA constructs against *Nanos1*, *Pum2*, *Cpsf4*, and *Smarcd1* were obtained from Dharmacon as glycerol stocks (Nanos1:RMM4534-EG332397, Pum2:RMM4534-EG80913, Cpsf4:RMM4534-EG54188, Smarcd1: RMM4534-EG83797). The sequences for all shRNA constructs were as follows (“*” indicates sh1 and “^” indicates sh2):

*shNanos1*
TRCN0000096771 – TAGCGCAGCTTCTTGCTGGGC
TRCN0000096769 – AACTTGAGCAATCAAGGTGGG*
TRCN0000096772 – TAGTGCGCGCAGTTCCAACGC
TRCN0000096770 – TACTTGATGGTATGTGCGTTG^
TRCN0000096773 – TGCAAACGGGTTCAGCTCAGC
*shPum2*
TRCN0000102260 – TTTGAACATGGTTAAAGCAGC*
TRCN0000102264 – ATGTCAGATCTATTATAGCGG
TRCN0000102261 – TTGCTGAAATAAATTGGCTGG
TRCN0000102262 – AAGAGTAGTAATATGAGGTCG^
TRCN0000102263 – ATCTTCCAATAACCTACTGCG
*shCpsf4*
TRCN0000123668 – ATGGGCAGTTCAAATCGAGGG
TRCN0000123666 – TATTCATGCAAGAACTCACAC*
TRCN0000123664 – AAATGAAAGGACAGACATGGC
TRCN0000123667 – AAATGCCCTTTGGTGCATCTG^
TRCN0000123665 – TTCACACAAATGACTCTCCGG
sh*Smarcd1*
TRCN0000092953 – TTTACCCGTTTGATTTCAGGC
TRCN0000092954 – AAGCTCAACATGAACTCTCGC*
TRCN0000092955 – TATGTGTTTCGGATTCCCAGG
TRCN0000092956 – ATCTCTGAGAACTTCATCCGC^
TRCN0000092957 – TACACCGTACATTCACATCTC

#### Overexpression

Mouse *Smarcd1* cDNA was purchased from Dharmacon as a glycerol stock (Accession: BC059921, Clone ID: 6816512, Catalog: MMM1013-202732694). Using directional TOPO-cloning primers (Fwd- 5’-CACCATGGCGGCCCGGGCGGGTTT-3’ and Rev-5’-TGTGTTTCGGATTCCCAGGGCTTGCTCTAACTCTTGCCGC-3’), *Smarcd1* cDNA was amplified using Phusion polymerase. DNA was purified using the QIAquick Gel Extraction Kit (Qiagen) and ligated into a Gateway entry clone using the pENTR/D-TOPO Cloning Kit (Invitrogen). Finally, pENTR-Smarcd1 was combined with pol2 promoter and C-terminal Myc-tag entry clones in a Gateway LR reaction according to the manufacturer’s protocol (Invitrogen).

### Virus transduction

1×10^6^ 293T cells were plated in 6 cm dishes 24 hours prior to transfection in P/S-free 10% FBS DMEM. Cells were transfected with 1 µg of shRNA/cDNA and 1 µg of viral packaging plasmids (250 ng pMD2.G and 750 ng psPAX2) using 6 µl of Xtreme Gene 9 transfection reagent (Roche). After 24 hours of transfection, media was refreshed with 10% DMEM, supplemented with 1% P/S and 1% glutamine. The following day, virus-containing supernatant was passed through a 45 µm filter to obtain viral particles, which were then transferred to 1×10^5^ 4T1/6DT1 cells. 24 hours post-transduction, the viral media was removed and fresh 10% DMEM was added. Finally, 48 hours after transduction, the cells were selected with 10 µg/ml puromycin- or 5 µg/ml blasticidin-containing complete DMEM.

### siRNA transfection

6DT1 cells were plated in P/S-free 10% FBS DMEM. 24 hours after plating, cells were transfected with ON-TARGETplus Non-targeting Control Pool siRNA or ON-TARGETplus Mouse Smarcd1 siRNA SMARTPool using DharmaFECT 1 Transfection Reagent (Dharmacon Inc, Horizon).

### Western blot

Protein lysates from 1×10^6^ cells were extracted on ice using Golden Lysis Buffer (10 mM Tris pH 8.0, 400 mM NaCl, 1% Triton X-100, 10% glycerol + Complete protease inhibitor cocktail (Roche), phosphatase inhibitor (Sigma)). Protein concentration was measured using Pierce’s BCA Protein Assay Kit and analyzed on a Versamax spectrophotometer at a wavelength of 560 nm. Appropriate volumes containing 20 µg of protein combined with NuPage LDS Sample Buffer and NuPage Reducing Agent (Invitrogen) were run on 4–12% NuPage Bis-Tris gels in MOPS buffer. Proteins were transferred onto a PVDF membrane (Millipore), blocked in 5% milk (dry milk diluted in Tris-buffered saline containing 0.05% Tween-20, TBST) for one hour and incubated in the primary antibody (in 5% milk) overnight at 4°C. Membranes were washed with TBST and secondary antibody incubations were done at room temperature for one hour. Proteins were visualized using the Amersham ECL Prime Western Blotting Detection System and Amersham Hyperfilm ECL (GE Healthcare). The following primary antibodies were used: mouse anti-CPSF4 (1:100; Santa Cruz), mouse anti-Pumilio2 (1:1,000; Abcam), mouse anti-SMARCD1 (1:1000; Bethyl A301-595A), rabbit anti-BICRA (1:1000; Abcam ab302712), rabbit anti-SMARCA4 (1:1000; Abcam ab110641), mouse anti-Actin (1:10,000; Abcam), mouse anti-Myc-tag (1:1000; Cell Signaling). Goat anti-rabbit (Santa Cruz) and goat-anti-mouse (GE Healthcare) secondary antibodies were used at concentrations of 1:10,000.

### RNA isolation, reverse transcription, and quantitative polymerase chain reaction

RNA was isolated from cell lines using TriPure (Roche) and reverse transcribed (RT) using iScript (Bio-Rad). Quantitative PCR of RT products (qRT-PCR) was conducted using VeriQuest SYBR Green qPCR Master Mix (Affymetrix). Peptidylprolyl isomerase B (*Ppib*) was used for normalization of expression levels. Expression of mRNA was defined from the threshold cycle, and relative expression levels were calculated using 2-delta Ct after normalization with *Ppib*. Primer sequences can be found in Table S11.

### Actinomycin D time course

Cellular *E2f2*, *Prr9*, and *Smarcd1* mRNA decay rates were measured using actinomycin D (actD) time course assays. Briefly, transcription was inhibited by addition of actD (5 µg/ml; Calbiochem) to the culture medium, and total RNA was purified at selected times thereafter. Time courses were limited to 4 hours to avoid complicating cellular mRNA decay pathways by actD-enhanced apoptosis^36^. mRNA levels were measured at each time point by qRT-PCR as described above. First-order decay constants (k) were solved by nonlinear regression (GraphPad Prism 9, San Diego, CA, USA) of the percentage of mRNA remaining versus time of actD treatment. Resolved mRNA half-lives (t1/2 = ln2/k) are based on the mean ± SD.

### *In vivo* metastasis assays

Female virgin FVB/NJ or BALB/cJ mice were obtained from The Jackson Laboratory at 6–8 weeks of age. Two days prior to *in vivo* experiments, cells were plated at 1×10^6^ cells per condition into T-75 flasks (Corning) in non-selective DMEM. Each mouse was injected with 1×10^5^ cells into the fourth mammary fat pad (orthotopic injection) or tail vein (tail vein injection). Recipient mice were of the FVB/NJ or BALB/cJ strains for 6DT1 or 4T1 cells, respectively. The mice were euthanized between 28–30 days post-injection. Primary tumors were resected, weighed, and lung metastases were counted.

### *In vitro* cellular phenotypic assays

#### Incucyte cell viability assays

Cells were seeded into 96-well plates at a density of 1×10^3^ cells per well and at least 4 wells per condition. Treatment was applied to cells 24 hours after seeding and then the plates were placed in a Sartorius Incucyte SX5 Live-Cell Analysis instrument in a 37°C incubator at 5% CO_2_, 20% O_2_. Cells were imaged every 4 hours using the SX5 G/R Optical Module phase channel at 10× magnification and analyzed with Incucyte 2022A Software for percent confluence in 4 regions per well. *Low glutamine:* media was replaced with DMEM + 10% FBS minus glutamine or with glutamine at 4 wells per condition for each cell line. *High ROS:* media was replaced with DMEM + 10% FBS or DMEM + 10% FBS + 500 µM H_2_O_2_ at 4 wells per condition for each cell line. *Heat shock:* Cells seeded in two 96-well plates, with one plate placed at 42°C 5% CO_2_ for 3 hours and the other kept at 37°C before placement of both into the Incucyte SX5. *Hypoxia:* Cells seeded in two 96-well plates, with one plate placed at 37°C at 5% CO_2_ and 1% O_2_ for 16 hours and the other kept at 20% O_2_ before placement of both into the Incucyte SX5 37°C at 5% CO_2_ and 20% O_2_. *Dox sensitivity:* Media was replaced with serial dilutions of Doxorubicin at concentrations between 100 to 0.001 nM in DMEM.

#### Colony assays

Cells were seeded in DMEM + 10% FBS into 6 wells of a 6-well plate at 6 serial dilutions: 1×10^4^, 5×10^3^, 1×10^3^, 5×10^2^, 1×10^2^, and 5×10^1^. After 5 days, media was removed and cells were stained using 1 ml crystal violet in 20% methanol for 10 minutes at room temperature before washing away excess stain with deionized water.

#### Viability under heat shock

Cells were seeded into a 96-well, flat clear bottom plates (Ibidi) at 5×10^3^ cells per well, 4 well replicates per cell line. After 24 hours, the media was replaced with DMEM + 10% FBS + CellEvent Caspase-3/7 Detection Reagent (Green) according to the manufacturers recommendations (Invitrogen). The plate was then imaged every 15 minutes using phase and GFP channels in the ZEISS Celldiscoverer 7 Automated Live Cell Imager set to 42°C at 5% CO_2_ and 20% O_2_. After 24 hours the images were assessed manually to determine average time until appearance of the first apoptotic cell (first GFP-positive cell) and average time to total cell death (all cells GFP positive) per field.

#### Cell migration scratch assay

Cells were seeded onto 35 mm dishes (Ibidi) and grown to 80% confluence. After 24 hours, two scratches were created in the cell monolayer of each dish using a pipet tip. The dishes were then washed several times to remove all dead and floating cells. Finally, DMEM + 10% FBS was added to the cells and the dishes were placed in the ZEISS Celldiscoverer 7 Automated Live Cell Imager set to 37°C at 5% CO_2_ and 20% O_2_. Scratches were imaged using the phase channel at three positions per scratch every three hours.

#### Tumorsphere assay

24-well ultralow attachment plates (Corning #3473) were prepared with 500 µl of a 1:1 ratio of MethoCult H4100 (Stemcell Technologies Cat# 04100) and complete MammoCult Media (MammoCult (StemCell Technologies Cat #05620) supplemented with 10% MammoCult Proliferation Supplement (StemCell Technologies Cat#05622), 1% hydrocortisone stock solution (StemCell Technologies Cat#07925), and 0.04% heparin (StemCell Technologies Cat #07980). Cells were then seeded at 5×10^3^ cells/well in 100 µl of complete MammoCult Media. For siRNA studies, the cells were seeded 24 hours after siRNA transfection. After culturing for 10 days, tumorspheres were imaged, sized, and quantitated using a Celigo Imaging Cytometer (Nexcelcom).

### RNA sequencing

#### Isolation of high-quality RNA

RNA was extracted from small sections of primary tumor tissue using TriPure, followed by organic extraction with chloroform and precipitation by isopropanol. RNA was then purified using the RNeasy Mini Kit (Qiagen) with on-column DNase digestion. RNA quality was tested using the Agilent 2200 TapeStation electrophoresis system, and samples with an RNA integrity number (RIN) score >7 were sent to the Sequencing Facility at Frederick National Laboratory.

#### Sequencing

Preparation of mRNA libraries and mRNA sequencing was performed by the Sequencing Facility using the HiSeq2500 instrument with Illumina TruSeq v4 chemistry.

#### Bioinformatic analysis

Sample reads were trimmed to remove adapters and low-quality bases using Trimmomatic software and aligned with the reference mouse mm9 genome and Ensemble v70 transcripts using Tophat software. RNA mapping statistics were calculated using Picard software. Library complexity was measured by unique fragments in the mapped reads using Picard’s MarkDuplicate utility. Gene-specific and transcript-specific analysis and differential expression analysis was performed using the Partek Genomics Suite. Genes with a p value 0.05 and false discovery rate (FDR) less than 0.05 were considered differentially expressed.

### Pathway analysis

#### Pathway Analysis (IPA)

Differentially expressed gene sets were analyzed using Ingenuity Pathway Analysis Software (Qiagen). Gene sets were uploaded into IPA for Core Expression Analysis of expression data. The Ingenuity Knowledge Base was chosen as the reference set of genes, and both direct and indirect relationships were considered. No other analysis parameters were specified, and the default settings were selected.

#### DAVID

A list of official gene symbols was uploaded into the DAVID Bioinformatics Resources 6.8 (https://david.ncifcrf.gov/) Functional Annotation Tool. Mus musculus was selected as the species and functional annotation clustering was selected using the default settings for annotation categories. Classification stringency was also left at the default setting of medium.

### Assays of transposase-accessible chromatin (ATAC)-seq library preparation

ATAC were performed as previously described^37^. Briefly, 5×10^4^ cells were isolated, and nuclei were generated by incubating on ice with 500 μl lysis buffer (RSB with 0.1% Tween-20) for 10 min. The resulting nuclei were centrifuged at 500 × g for 10 min, then resuspended in 1X Tagment DNA buffer (Illumina) with 2.5 μl Tagment DNA Enzyme (Illumina) and incubated at 37 °C for 30 min. For each transposition reaction, the volume was 50 μl. The transposition mixtures were quenched with 500 μl PB buffer (Qiagen) and purified by standard protocol with the MinElute PCR purification kit. Each ATAC library was amplified with Nextera primers for 16 PCR cycles and purified with Agencourt AMPure XP (Beckman Coulter) to remove excess primers. The resulting ATAC libraries were sequenced with NextSeq500 with paired-end reads.

### Analysis of ATAC-seq data

In Partek Flow, ATAC-seq reads were aligned to the mouse genome version mm39 using Bowtie2 (Version 2.2.5) and unaligned reads filtered out. Peak calling was then performed using MACS (version 3.00a7) and annotated with Ensemble Transcripts (release 109 prajapatmk), specifying 1 gene region per peak. Finally, known motif detection was carried out with the All CORE profiles database and *de novo* motif detection using a length of 6-16 base pairs.

### Co-immunoprecipitation, silver staining, and proteomics analysis

Co-immunoprecipitation (co-IP) was performed using the Nuclear Complex Co-IP Kit (Active Motif). 6T1 *Smarcd1* OE cells were seeded onto 15 cm tissue culture dishes at a seeding density of 5×10^6^ cells per dish. After 48 hours of incubation, cells were harvested, and nuclear lysates were prepared. A total of 200–500 μg of nuclear lysates were incubated with 2 μg of SMARCD1 antibody (Bethyl A301-594A) and 50 μg of Dynabeads Protein G (Invitrogen). After overnight incubation on a rotator at 4 °C, immune complexes were isolated using a magnetic stand. Beads were then washed three times, resuspended in 2× NuPAGE LDS sample buffer (Invitrogen), and incubated at 95 °C in a heat block for 5 min. Samples were loaded onto NuPAGE protein gels and the standard western blot protocol was followed as described above. For silver stain analysis, the NuPAGE protein gel was processed and stained using Pierce Silver Stain for Mass Spectrometry (Thermo Fisher). For tandem Mass Spectrometry (MS/MS) and proteomics analysis, the co-IP samples still on the beads were sent to the NCI Collaborative Protein Technology Resource. Briefly, samples were solution-digested with trypsin using S traps (Protifi), following the manufacturer’s instructions. The digested peptides were analyzed on an Orbitrap Exploris 480 (Thermo) mass spectrometer inline after an UltiMate 3000 RSLCnano HPLC (Thermo). The peptides were separated on a 75 µm x 15 cm, 3 µm Acclaim PepMap reverse phase column (Thermo) and eluted directly into the mass spectrometer. For analysis, parent full-scan mass spectra were acquired at 120,000 FWHM resolution and product ion spectra at 15,000 resolution. Proteome Discoverer 3.0 (Thermo) was used to search the data against the murine database from Uniprot using SequestHT with INFERYS rescoring (Zolg et al, PMID: 34015160). The search was limited to tryptic peptides, with maximally two missed cleavages allowed. Cysteine carbamidomethylation was set as a fixed modification, with methionine oxidation as a variable modification. The precursor mass tolerance was 10 ppm, and the fragment mass tolerance was 0.02 Da. The Percolator node was used to score and rank peptide matches using a 1% false discovery rate. Label-free quantitation of extracted ion chromatograms from MS1 spectra was performed using the Minora node; missing values were replaced with 100 as a minimum quantitation threshold.

### Ethics statement

The research described in this study was performed under the Animal Study Protocol LCBG-004, approved by the NCI Bethesda Animal Use and Care Committee. Animal euthanasia was performed by cervical dislocation after anesthesia by Avertin.

### Patient data

Kaplan-Meier analysis of the *CPSF4*, *PUM2*, and *NANOS1* gene list as a signature was performed using Kaplan-Meier plotter (https://kmplot.com) and breast cancer data sets. The genes were weighted equally, and T1 and T3 of trichotomized DMFS curves using mean expression in tumor tissue were generated for the PAM50 basal (n=1671) patient cohort. Kaplan-Meier DMFS analysis for all other genes and gene list signatures was performed using Gene Set analysis – Tumors tool within The Gene expression-based Outcome for Breast cancer Online (GOBO) site (http://co.bmc.lu.se/gobo/).

### Data analysis and statistics

Comparisons of mRNA levels and decay kinetics were performed using the unpaired t test. Differences yielding p < 0.05 were considered significant. Statistical significance between groups in *in vivo* assays was determined using the Mann-Whitney unpaired nonparametric test using Prism (version 5.03, GraphPad Software, La Jolla, CA). Correlation analyses used the Spearman non-parametric test while Kaplan-Meier comparisons were performed using the log-rank test with events limited to death from recurrent disease. For correlation and survival analyses, differences yielding p < 0.05 were considered significant.

## Acknowledgements

This research was supported by the Intramural Research Program of the NIH. We are grateful to the members of the NCI Collaborative Protein Technology Resource for their help with mass spectrometry and data analysis; to the CCR animal resource center for supporting our *in vivo* experiments; The CCR Sequencing Facility, as well as the Kahn laboratory for guidance with ATAC-seq protocol development. We also thank the members of the Laboratory of Cancer Biology and Genetics for experimental ideation and project discussion, and specifically Meera Murgai PhD and Brandi Carofino PhD for their help with manuscript preparations.

