## Supplemental Table Legends for "SMARCD1 is a “Goldilocks” metastasis modifier"

**Supplemental Tables**

**Table S1: NANOS1-, PUM2-, and CPSF4-responsive genes and DAVID analysis.**

Sheet 1: NANOS1-responsive genes. Differential gene expression analysis compared to control.

Sheet 2: PUM2-responsive genes. Differential gene expression analysis compared to control.

Sheet 3: CPSF4-responsive genes. Differential gene expression analysis compared to control.

Sheet 4: Venn diagram analysis. 21 genes commonly altered in all knockdown lines compared to control.

Sheet 5: DAVID functional clustering analysis of genes listed in sheet 4.

**Table S2: Ingenuity Pathway Analysis of differential gene expression in NANOS1, PUM2, and CPSF4 knockdown (KD) cells vs control.**

Sheet1: IPA analysis of differential gene expression in NANOS1 KD cells compared to control.

Sheet2: IPA analysis of differential gene expression in PUM2 KD cells compared to control.

Sheet3: IPA analysis of differential gene expression in CPSF4 KD cells compared to control.

**Table S3: *E2f2*, *Prr9*, and *Smarcd1* mRNA half-life summary table.**

**Table S4: SMARCD1-responsive genes in monolayer and 3D culture.**

Sheet 1: Differential gene expression analysis for *Smarcd1* overexpression (OE) vs empty vector (EV) control cells grown in monolayer.

Sheet 2: Differential gene expression analysis for *Smarcd1* knockdown (KD) vs shScr control cells grown in monolayer.

Sheet 3: Venn diagram analysis of differentially expressed genes in *Smarcd1* OE vs EV cells and KD vs shScr cells grown in monolayer.

Sheet 4: Differential gene expression analysis for *Smarcd1* OE vs EV cells grown in 3D.

Sheet 5: Differential gene expression analysis for *Smarcd1* KD vs shScr cells grown in 3D.

Sheet 6: Venn diagram analysis of differentially expressed genes in *Smarcd1* OE vs EV cells and KD vs shScr cells grown in 3D.

**Table S5: ATAC-seq annotated peaks.**

Sheet 1: Annotated regions of open chromatin in 6DT1 empty vector (EV) cells grown in monolayer.

Sheet 2: Annotated regions of open chromatin in 6DT1 shScramble (shScr) cells grown in monolayer.

Sheet 3: Annotated regions of open chromatin in 6DT1 *Smarcd1* overexpression (OE) cells grown in monolayer.

Sheet 4: Annotated regions of open chromatin in 6DT1 *Smarcd1* knockdown (KD) cells grown in monolayer.

Sheet 5: Annotated regions of open chromatin in 6DT1 EV cells grown in 3D (spheres).

Sheet 6: Annotated regions of open chromatin in 6DT1 shScr cells grown in 3D (spheres).

Sheet 7: Annotated regions of open chromatin in 6DT1 Smarcd1 OE cells grown in 3D (spheres).

Sheet 8: Annotated regions of open chromatin in 6DT1 Smarcd1 KD cells grown in 3D (spheres).

**Table S6: Unique motif enrichment in open chromatin.**

Sheet 1: Venn diagram analysis identifying significantly enriched motifs unique to each cell line.

Sheet 2: Significantly enriched motifs within open chromatin of 6DT1 empty vector (EV) cells grown in monolayer.

Sheet 3: Significantly enriched motifs within open chromatin of 6DT1 shScramble (shScr) cells grown in monolayer.

Sheet 4: Significantly enriched motifs within open chromatin of 6DT1 *Smarcd1* overexpression (OE) cells grown in monolayer.

Sheet 5: Significantly enriched motifs within open chromatin of 6DT1 *Smarcd1* knockdown (KD) cells grown in monolayer.

Sheet 6: Significantly enriched motifs within open chromatin of 6DT1 EV cells grown in 3D (spheres).

Sheet 7: Significantly enriched motifs within open chromatin of 6DT1 shScr cells grown in 3D (spheres).

Sheet 8: Significantly enriched motifs within open chromatin of 6DT1 Smarcd1 OE cells grown in 3D (spheres).

Sheet 9: Significantly enriched motifs within open chromatin of 6DT1 Smarcd1 KD cells grown in 3D (spheres).

**Table S7: SMARCD1-interacting proteins.**

**Table S8: Differential splicing in *Smarcd1*-altered cells.**

Sheet 1: Differential splicing analysis for *Smarcd1* overexpression (OE) vs empty vector (EV) cells grown in monolayer.

Sheet 2: Differential splicing analysis for *Smarcd1* knockdown (KD) vs shScramble (shScr) cells grown in monolayer.

Sheet 3: Venn diagram analysis of differentially spliced transcripts in *Smarcd1* OE vs EV cells and KD vs shScr grown in monolayer.

Sheet 4: Differential splicing analysis for *Smarcd1* OE vs EV cells grown in 3D.

Sheet 5: Differential splicing expression analysis for *Smarcd1* KD vs shScr cells grown in 3D.

Sheet 6: Venn diagram analysis of differentially spliced transcripts in *Smarcd1* OE vs EV cells and KD vs shScr grown in 3D.

**Table S9: Ingenuity Pathway Analysis (IPA) of differentially spliced transcripts genes in *Smarcd1*-altered cells vs controls.**

Sheet 1: IPA of differentially spliced genes in *Smarcd1* overexpression (OE) cells compared to control grown in monolayer.

Sheet 2: IPA of differentially spliced genes in *Smarcd1* knockdown (KD) cells compared to control grown in monolayer.

Sheet 3: IPA of differentially spliced genes in *Smarcd1* OE cells compared to control cells grown in 3D (spheres).

Sheet 4: IPA of differentially spliced genes in *Smarcd1* KD cells compared to control cells grown in 3D (spheres).

Sheet 5: Venn diagram analysis for monolayer-specific pathways enriched in alternate splice variants, categorized by cellular process.

**Table S10: Manual assessment of key cellular machinery for a “Goldilocks pattern” of distant metastasis-free survival (DMFS) using the GOBO dataset and KM analysis.**

**Table S11: qRT-PCR primer sequences.**
