## Supplemental Figures S1-S8 for "SMARCD1 is a “Goldilocks” metastasis modifier"

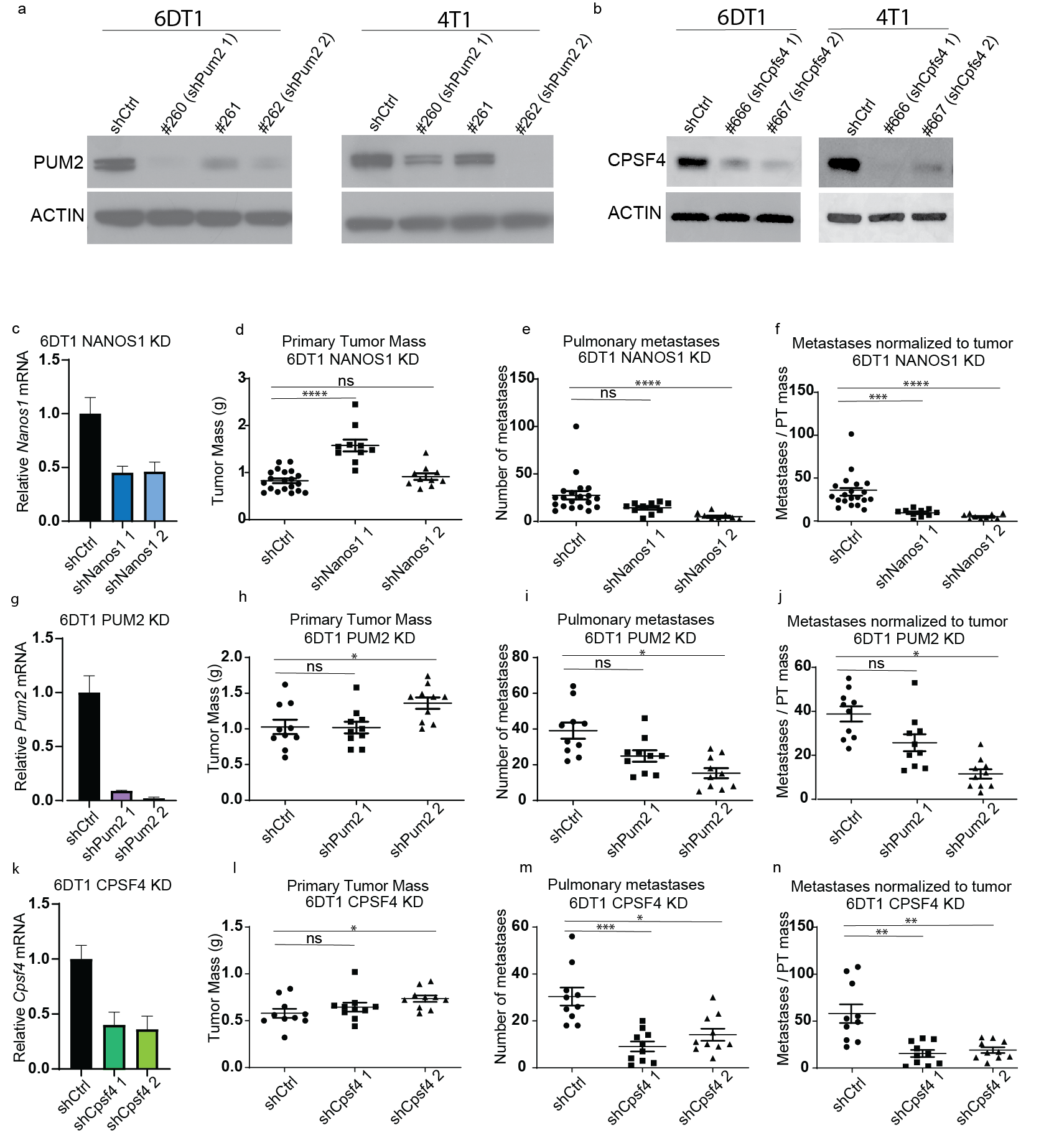

**Figure S1. RNA-binding proteins NANOS1, PUM2, and CPSF4 are metastasis modifiers in 6DT1 mouse mammary breast cancer cells.**

Figure S1a, western blot analysis of PUM2 protein levels in 6DT1 and 4T1 cells transduced with scramble shScr or three different shRNAs targeted at *Pum2* mRNA. b, western blot analysis of CPSF4 protein levels in 6DT1 and 4T1 cells transduced with scramble shScr or three different shRNAs targeted at *Cpsf4* mRNA. c, qRT-PCR analysis of *Nanos1* expression in 6DT1 cells transduced with shScr, or shNanos1 1 or 2. d-f, analysis of primary tumor (PT) weight (d), number of lung nodules (e), and number of lung nodules per gram of PT per mouse (f) 28 days after orthotopic injection of 6DT1 shScr or shNanos1 KD lines. g, qRT-PCR analysis of *Pum2* expression in 6DT1 cells transduced with shScr, or shPum2 1 or 2. h-j, analysis of PT weight (h), number of lung nodules (i), and number of lung nodules per gram of PT per mouse (j) 28 days after orthotopic injection of 6DT1 shScr or shPum2 KD lines. k, qRT-PCR analysis of *Cpsf4* expression in 6DT1 cells transduced with shScr, or shCpsf4 1 or 2. l-n, analysis of PT weight (l), number of lung nodules (m), and number of lung nodules per gram of PT per mouse (n) 28 days after orthotopic injection of 6DT1 shScr or shCpsf4 KD lines. n.s = not statistically significant, * = p<0.05, ** = p<0.01, ***=p<0.001, ****=p<0.0001.

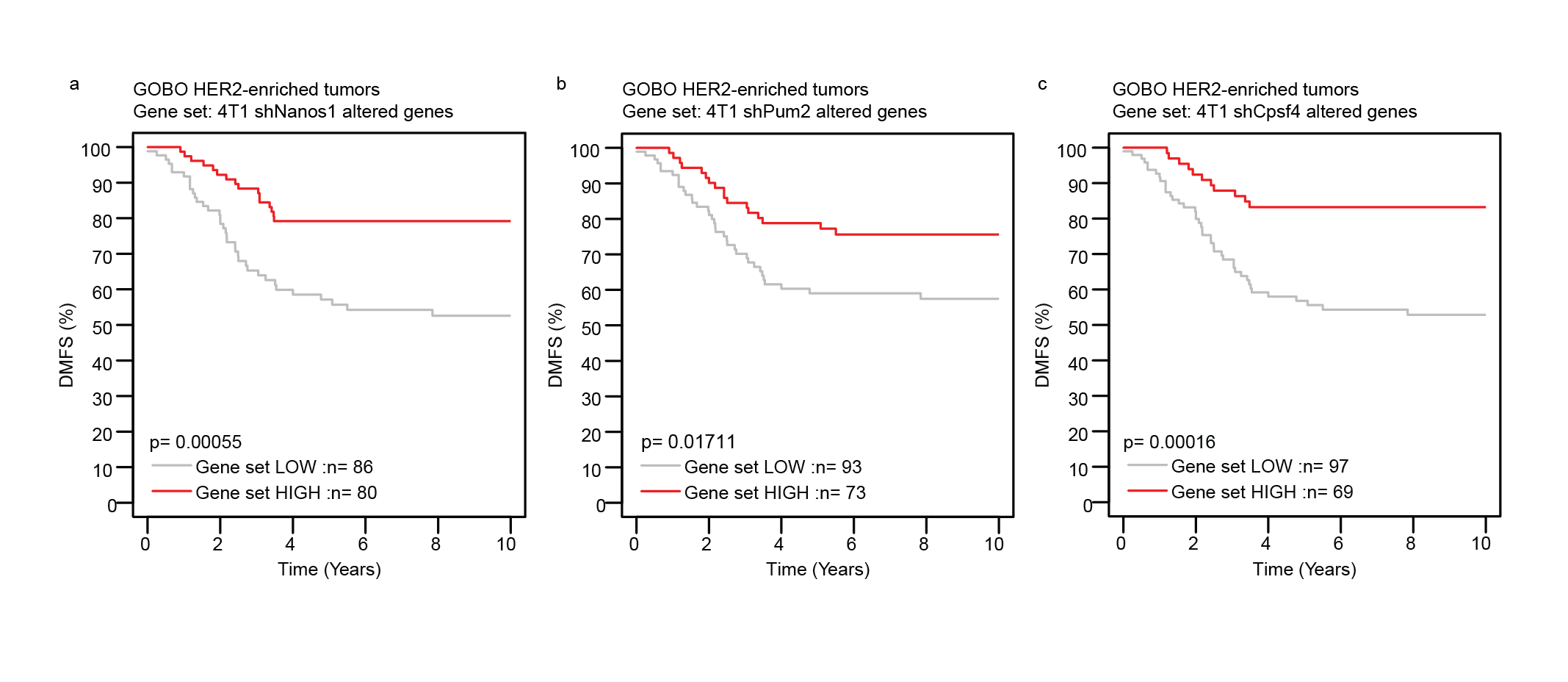

**Figure S2. Differentially expressed genes from RNA-binding protein knockdown lines stratify ER+/HER2-enriched patient distant metastasis-free survival.**

Figure S2a-c, Kaplan-Meier analysis of distant metastasis-free survival (DMFS) for ER+/HER2-enriched patients stratified by differentially expressed gene lists from 4T1 shNanos1 cells (a), shPum2 cells (b), and shCpsf4 cells (c) used as non-weighted signatures in the GOBO dataset.

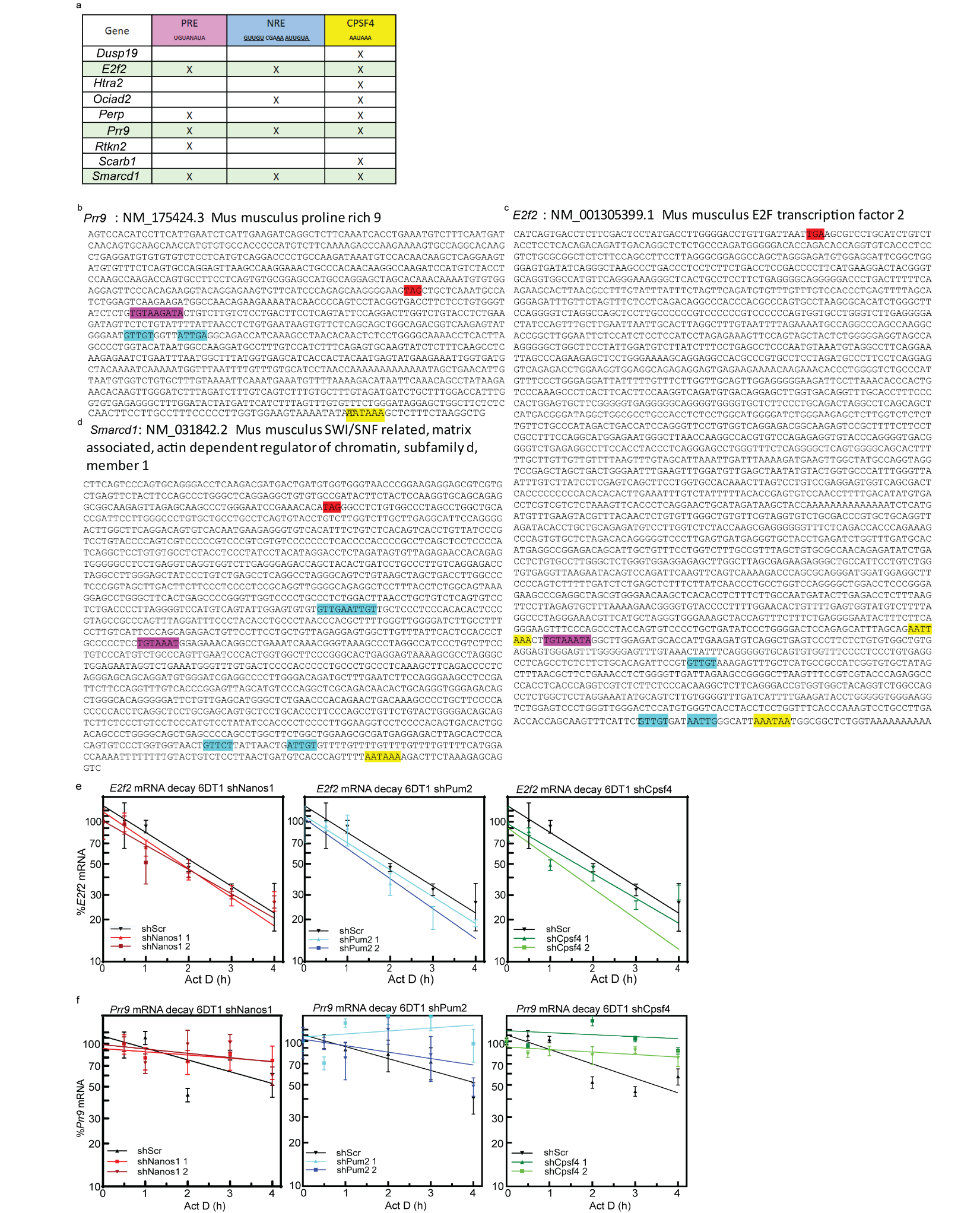

**Figure S3. Sequence and half-life analysis of differentially expressed transcripts in RNA-binding protein knockdown lines.**

Figure S3a, table showing the presence of a PRE, NRE, or CPSF4 binding elements within the 3’UTR of each gene listed where x = presence of element. b-d, 3’UTR sequences with PRE, NRE, CPSF4 binding elements highlighted (red: stop codon, purple: PRE, blue: NRE, yellow: CPSF4 binding element) in the *Prr9* 3’UTR (b), *E2f2* 3’UTR (c), and *Smarcd1* 3’UTR (d). e, graphs showing actinomycin D (actD) time course to measure *E2f2* RNA half-life in 6DT1 shNanos1 (red), shPum2 (blue), shCpsf4 (green) vs shScr. f, graphs showing actD time course to measure *Prr9* RNA half-life in 6DT1 shNanos1 (red), shPum2 (blue), shCpsf4 (green) vs shScr.

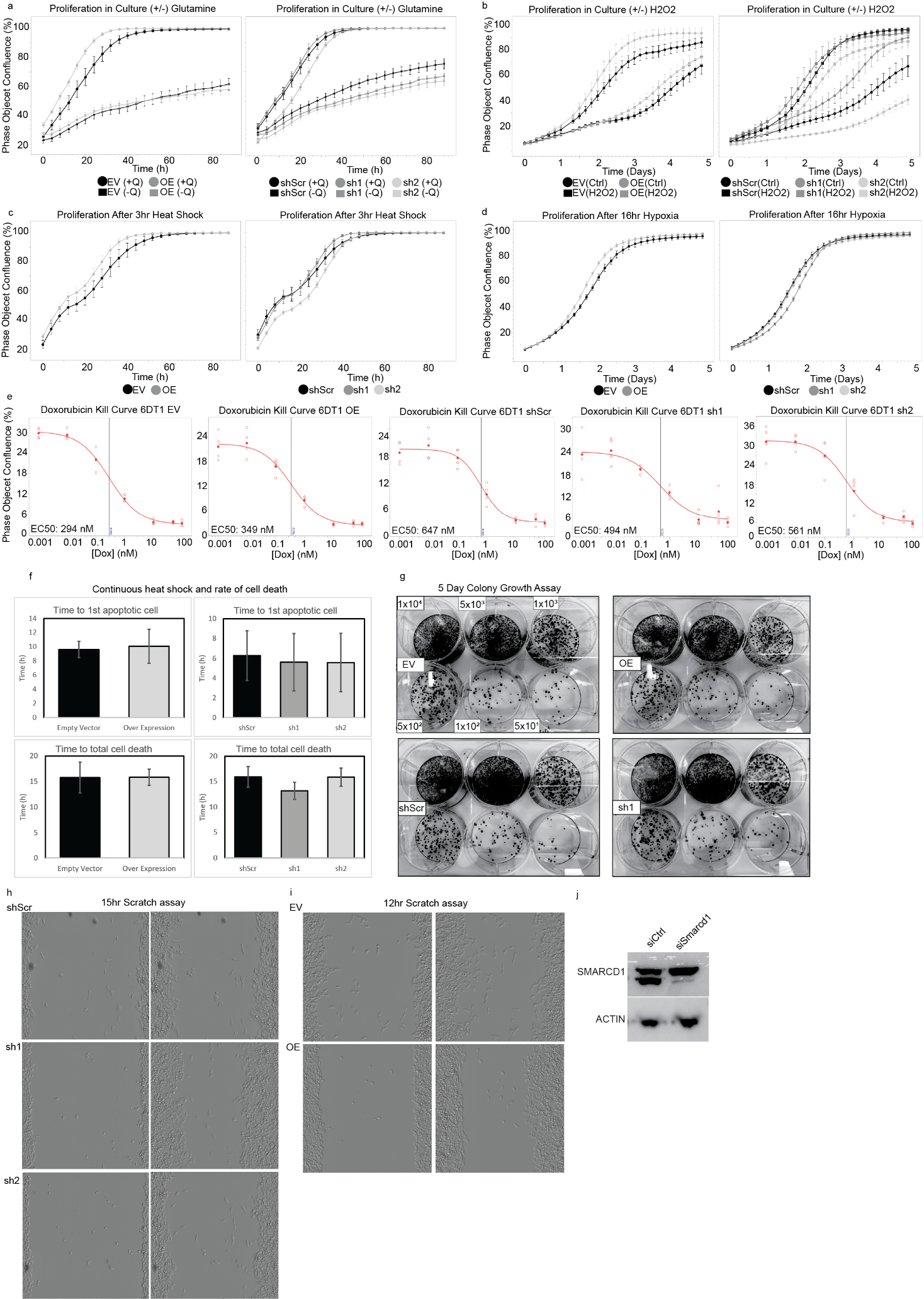

**Figure S4. Altered *Smarcd1* expression does not impact cell response to stress.**

Figure S4a-d, graphs showing phase object cell confluence over time following treatment with “no glutamine” media (a), treatment with H_2_O_2_ (b), 3 hour heat shock at 42°C (c), and 16 hours in hypoxic 1% oxygen conditions (d) (n=4). e, kill curves showing treatment curves and EC50 values for each cell line after 24 hour treatment with a serial dilution series of doxorubicin (n=4). f, histograms showing time to appearance of the first apoptotic cell and time until all cells were apoptotic during 20 hours of continuous heat shock (n=10). g, photographs of colony growth assays with cells seeded in 6-well plates at limiting dilutions. h, representative images of migration assays at time point 0 and at 15 hours in 6DT1 *Smarcd1* KD lines vs shScr control. i, representative images of migration assays at time point 0 and 12 hours in 6DT1 *Smarcd1* OE lines vs EV control. j, western blot showing SMARCD1 and Actin protein levels after transfections with control “siCtrl” or *Smarcd1*-targeted “siSmarcd1” siRNA.

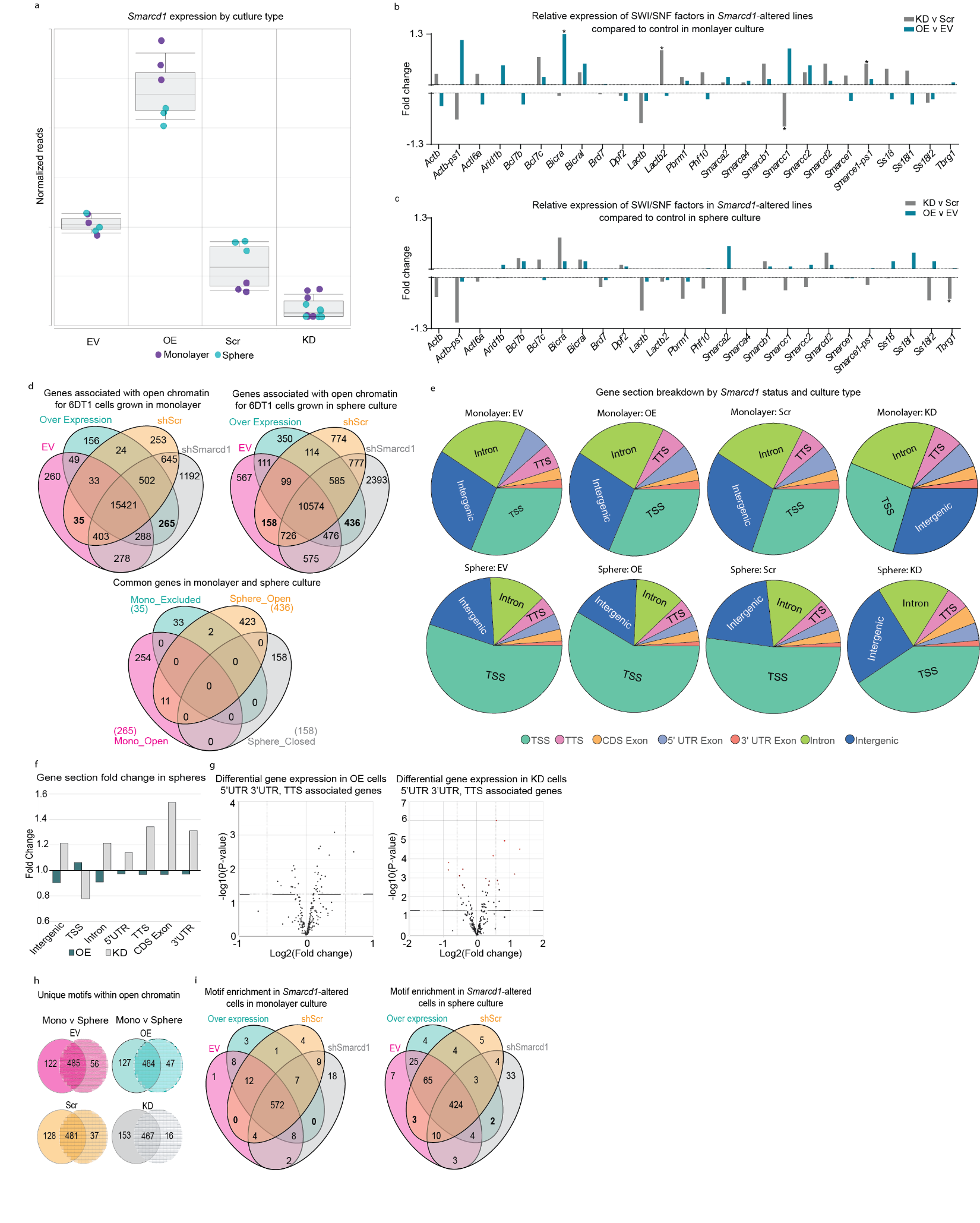

**Figure S5. Impact of altered *Smarcd1* expression on gene expression and chromatin accessibility.**

Figure S5a, RNA-seq data showing *Smarcd1* mRNA levels in 6DT1 EV, OE, Scr, and KD lines in monolayer and sphere culture. b-c, RNA-seq data showing relative abundance SWI/SNF factor mRNA in *Smarcd1*-altered cells vs controls in monolayer (b) and sphere culture (c). d, Venn diagram analysis of genes associated with regions of open chromatin for each cell line according to culture type. e, pie charts showing the relative percentage of each gene section found in sequenced open chromatin. f, graph showing fold change in the enrichment of specific gene sections (Intergenic, Transcription start site or TSS, Intron, 5 prime untranslated region or 5’UTR, Transcription termination sequence or TTS, coding sequence or CDS, and 3’UTR) in open chromatin for *Smarcd1*-altered cells vs control lines grown as spheres. g, volcano plots showing changes in gene expression for genes with 3’UTR, 5’UTR, and TTS enrichment in open chromatin in monolayer culture (red=fold change >1.5, blue=fold change <1.5). h, Venn diagram analysis of unique motifs enriched in open chromatin for each cell line in monolayer vs sphere culture. i, Venn diagram analysis of motifs enriched in open chromatin for each cell line in monolayer or sphere culture.

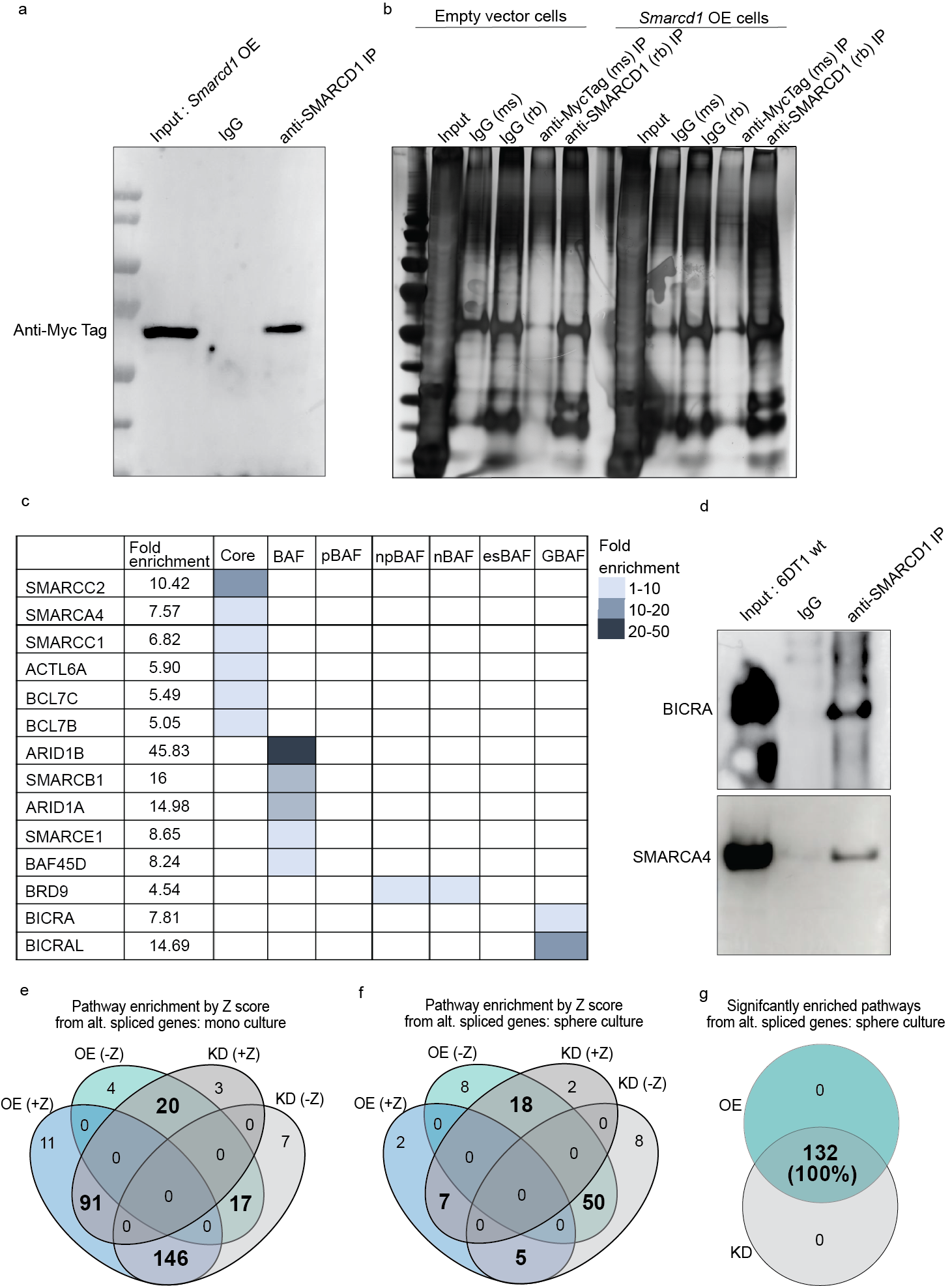

**Figure S6. SMARCD1 interacts with BAF complex members and alters splicing programs in sphere culture.**

Figure S6a, western blot showing efficiency and specificity of SMARCD1 immunoprecipitation (IP) from 6DT1 *Smarcd1* overexpression (OE) cells. b, silver-stained gel showing relative protein abundance in input, IgG control, Myc-tag, and SMARCD1 antibody IP samples. c, table showing fold enrichment of BAF complex members in *Smarcd1* OE vs EV. d, western blot showing that BICRA and SMARCD4 immunoprecipitate with SMARCD1. e-f, Venn diagram analysis of pathway analysis Z scores in *Smarcd1* OE and KD lines vs controls in monolayter “mono” culture (e) and sphere culture (f). g, Venn diagram analysis of pathways significantly enriched (p<0.05) by alternatively spliced genes in OE and KD lines from sphere culture.

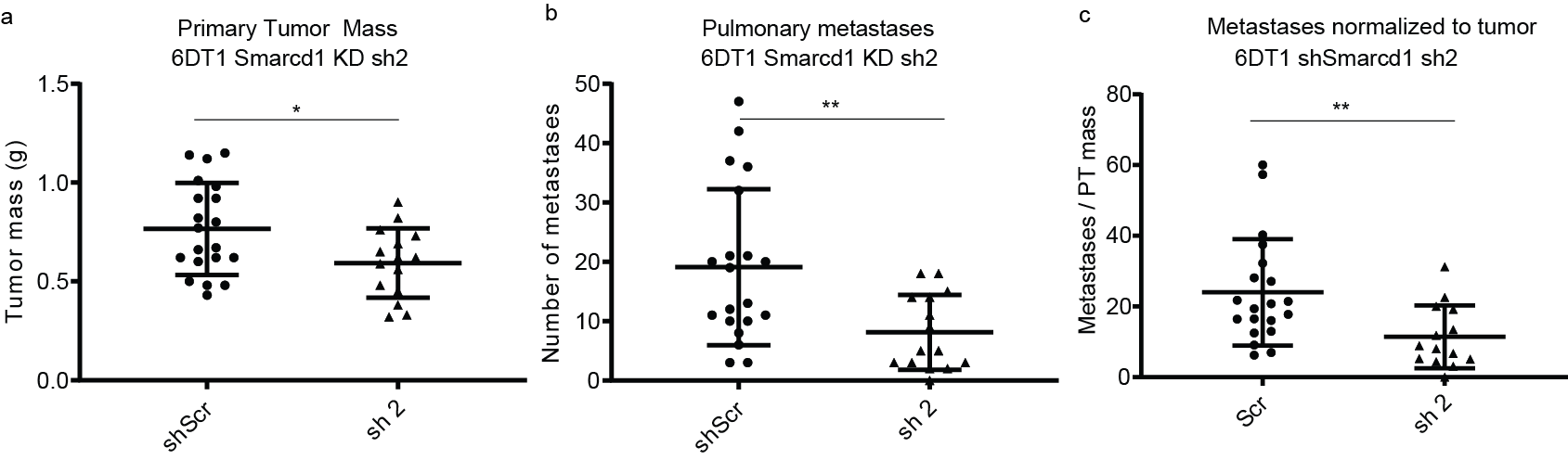

**Figure S7. *Smarcd1* KD decreases metastasis in an orthotopic mouse model.**

a-c, analysis of primary tumor (PT) weight (a), number of lung nodules (b), and number of lung nodules per gram of PT per mouse (c) 28 days after orthotopic injection of 6DT1 shScr control and shSmarcd1 line 2 (2 combined experiments, 10 mice per experiment).* = p<0.05, ** = p<0.01.

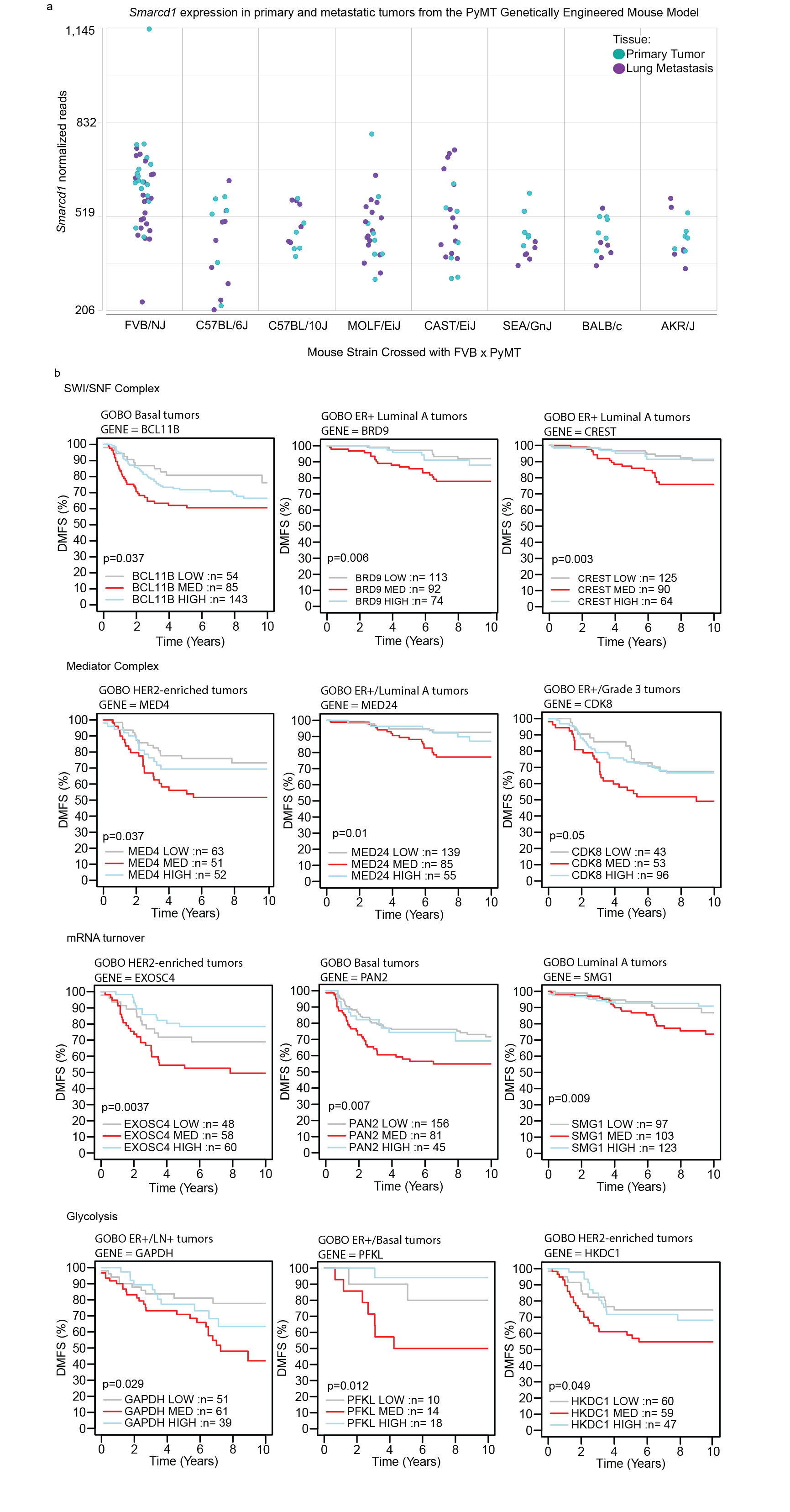

**Figure S8. Goldilocks metastasis modifier genes exist throughout cellular processes and clinical subtypes.**

Figure S8a, graph of *Smarcd1* RNA levels from RNA-seq data of matched primary tumor (PT) and lung metastasis pairs from the FVB x PyMT GEMM crossed to several mouse strains. Each point is data from 1 mouse, green=PT tissue, purple=metastatic tissue. b, representative Kaplan-Meier analysis plots showing stratification of distant metastasis-free survival (DMFS) for patients with different subtypes of breast cancer by expression level of the indicated genes (grey=low expression, red=medium expression, blue=high expression).
